# Cortical Parvalbumin-positive Interneuron Development and Function are Altered in the APC Conditional Knockout Mouse Model of Infantile Spasm Syndrome

**DOI:** 10.1101/2022.07.06.499046

**Authors:** Rachael F. Ryner, Isabel D. Derera, Moritz Armbruster, Anar Kansara, Mary E. Sommer, Antonella Pirone, Farzad Noubary, Michele Jacob, Chris G. Dulla

**Affiliations:** Department of Neuroscience, Tufts University School of Medicine, Boston, Massachusetts, USA.; Cell, Molecular, and Developmental Biology Graduate Program, Tufts Graduate School of Biomedical Sciences, Boston, Massachusetts, USA.; Department of Health Sciences, Bouvé College of Health Sciences, Northeastern University, Boston, Massachusetts, USA.

**Author notes:** Corresponding Author: Chris G. Dulla, Department of Neuroscience, Tufts University School of Medicine South Cove 203, 136 Harrison Avenue, Boston, Massachusetts 02111.

## Abstract

Infantile Spasms syndrome (ISS) is a childhood epilepsy syndrome characterized by infantile or late onset spasms, abnormal neonatal EEG, and epilepsy. Few treatments exist for IS, clinical outcomes are poor, and the molecular and circuit-level etiologies of IS are not well understood. Multiple human ISS risk genes are linked to Wnt/β-catenin signaling, a pathway which controls developmental transcriptional programs and promotes glutamatergic excitation via β-catenin’s role as a synaptic scaffold. We previously showed that deleting adenomatous polyposis coli (APC), a component of the β-catenin destruction complex, in excitatory neurons (APC cKO mice, APC^fl/fl^ x CaMKIIα^Cre^) in mice increased β-catenin levels in developing glutamatergic neurons and led to infantile behavioral spasms, abnormal neonatal EEG, and adult epilepsy. Here, we tested the hypothesis that the development of inhibitory GABAergic interneurons (INs) is disrupted in APC cKOs. IN dysfunction is implicated in human ISS, is a feature of other rodent models of ISS and may contribute to the manifestation of spasms and seizures. We found that parvalbumin positive INs (PV+INs), an important source of cortical inhibition, were decreased in number, underwent disproportionate developmental apoptosis, and had altered dendrite morphology at P9, the peak time of behavioral spasms. PV+INs received excessive excitatory input and their intrinsic ability to fire action potentials was reduced at all timepoints examined (P9, P14, P60). Subsequently, synaptic inhibition of pyramidal neurons was uniquely altered in the somatosensory cortex of APC cKO mice at all ages, with both decreased inhibition at P14 and enhanced inhibition at P9 and P60. These results indicate that inhibitory circuit dysfunction occurs in APC cKOs and, along with known changes in excitation, may contribute to ISS-related phenotypes.

**Significance Statement:** Infantile spasms syndrome (ISS) is a devastating epilepsy with limited treatment options and poor clinical outcomes. The molecular, cellular, and circuit disruptions that cause infantile spasms and seizures are largely unknown, but inhibitory GABAergic interneuron dysfunction has been implicated in rodent models of ISS and may contribute to human ISS. Here, we utilize a rodent model of ISS, the APC cKO mouse, in which β-catenin signaling is increased in excitatory neurons. This results in altered parvalbumin-positive GABAergic interneuron development and inhibitory synaptic dysfunction throughout life, showing that pathology arising in excitatory neurons can initiate long-term interneuron dysfunction. Our findings further implicate GABAergic dysfunction in ISS, even when pathology is initiated in other neuronal types.

## Introduction

Infantile Spasms Syndrome (ISS) is a devastating developmental epilepsy syndrome that occurs in the first year of life (West, 1841; Pavone et al., 2014) and is characterized by flexion/extension spasms, neurodevelopmental delays, and hypsarrhythmia (Paciorkowski et al., 2011; Michaud et al., 2014; Boutry-Kryza et al., 2015; Pirone et al., 2017). Early suppression of spasms and epileptiform activity can improve clinical outcomes (Shields, 2006; El Achkar and Spence, 2015), but available therapies are not widely effective and have adverse effects (Go et al., 2012; Riikonen et al., 2015). The pathological processes that lead to ISS are not well defined, but inhibitory GABAergic network dysfunction is implicated across multiple ISS mouse models (Marsh et al., 2009; Price et al., 2009; Olivetti et al., 2014; Marsh et al., 2016; Katsarou et al., 2018). In addition, GABAergic dysfunction in seen in a mouse model of TSC (Fu et al., 2012), which in humans is associated with ISS (note: the TSC mouse model has not been demonstrated to have spasms). Finally, GABAergic dysfunction occurs in human neurological disorders associated with ISS, some of which may be considered “interneuronopathies” (Bonneau et al., 2002; Kato and Dobyns, 2005). Understanding the mechanisms that lead to synaptic, cellular, and circuit dysfunction in ISS, and what differentiates spasms from seizures on a circuit-level, is essential to developing novel therapeutic interventions.

Interestingly, several genes linked to ISS, including Arx, LIS1, FOXG1, and TSC1/2 (Stromme et al., 2002a; Stromme et al., 2002b; Paciorkowski et al., 2011; Striano et al., 2011; Michaud et al., 2014) associate with the β-catenin signaling pathway. β-catenin is strongly implicated in neurodevelopmental disorders (Cotter et al., 1998; Zhang et al., 1998; Mohn et al., 2014a; Kuechler et al., 2015; Wickham et al., 2019; Alexander et al., 2020) and plays a dual role in 1) mediating transcriptional activation of Wnt-target genes (Behrens et al., 1996; Molenaar et al., 1996) and 2) as a structural element stabilizing synapses via its interactions with classic cadherins (Bamji et al., 2003; Kiryushko et al., 2004; Schlessinger et al., 2009; Salinas, 2012; Vitureira et al., 2012). Arx is expressed in GABAergic interneurons (INs) (Colombo et al., 2004; Poirier et al., 2004; Friocourt et al., 2006) and binds to β-catenin to positively regulate downstream signaling (Cho et al., 2017). The adenomatous polyposis coli (APC) protein, part of the β-catenin destruction complex, associates with FOXG1 and LIS1, regulating Wnt signaling and directing cell fate in the developing brain (Kato and Dobyns, 2003; McManus et al., 2004; Hebbar et al., 2008; Danesin et al., 2009; Ivaniutsin et al., 2009; Miyoshi and Fishell, 2012). TSC1/2 functions as a repressor of β-catenin and conditional deletion of TSC1/2 in GABAergic IN progenitors decreases seizure threshold (Mak et al., 2005; Huang and Manning, 2008; Carson et al., 2013). Although these data provide correlative evidence linking ISS to APC/Wnt/β-catenin signaling, there are few studies that directly link the two.

To determine the link between β-catenin and ISS, we utilized APC conditional homozygous knock- out mice (APC^fl/fl^ crossed with CaMKIIα^CRE^/APC^fl/+^, APC cKO) which lack APC in excitatory forebrain neurons beginning in early postnatal life, leading to elevated levels of β-catenin protein (Mohn et al., 2014a). APC cKO mice recapitulate phenotypes relevant to human ISS including infantile flexion/extension spasms, abnormal neonatal EEG activity, and spontaneous electroclinical seizure activity in adult mice (Pirone et al., 2017). APC cKO mice also show intellectual disabilities and behavioral abnormalities considered relevant to human autism spectrum disorder (Mohn et al., 2014a), which are often comorbid with ISS (El Achkar and Spence, 2015). Consistent with β-catenin’s role in synapse density and maturation, APC cKO mice show increased immature spine density and enhanced excitatory neurotransmission in the hippocampus and cortex as early as post-natal day 9 (P9), the same developmental time point of peak behavioral spasms in this model. Cortical circuit maturation is activity-dependent, therefore aberrant excitation in early life could drive long-term changes in brain function and lead to adult seizures. Of particular interest, the survival and maturation of GABAergic INs are dynamically regulated by excitatory activity during early development (Southwell et al., 2012; Priya et al., 2018; Wong et al., 2018). GABAergic interneurons (IN) provide inhibitory constraint of cortical circuits, are critical in developmental cortical circuit refinement (Freund and Buzsaki, 1996; Buzsáki, 2006; Klausberger and Somogyi, 2008), and GABAergic circuit dysfunction is implicated in epilepsy and ISS (Yu et al., 2006; Price et al., 2009; Khazipov, 2016). Interestingly, parvalbumin-positive (PV+) IN density has been reported to be decreased in the adult prefrontal cortex of APC cKO mice (Pirone et al., 2018), but the mechanisms by which PV+IN density decreases and how this affects inhibitory synaptic function is unknown.

We hypothesized that GABAergic IN survival and maturation may be altered in the APC cKO mouse model of ISS. β-catenin regulates both cortical excitation (Bamji et al., 2003; Wisniewska et al., 2012) and GABAergic IN development (Gulacsi and Anderson, 2008), providing rationale to suspect that GABAergic IN maturation may be disrupted in the APC cKO model of ISS. Here we used genetic, anatomical, and electrophysiological approaches to show that parvalbumin (PV+) INs, an important source of cortical inhibition, are reduced in number, undergo increased levels of developmental apoptosis, and receive excessive excitation in the somatosensory cortex of neonatal APC cKO mice. Consistent with early life excitation driving long-term changes in GABAergic IN function, adult APC cKOs show altered synaptic inhibition of excitatory cortical neurons and PV+IN intrinsic excitability. Overall, our study shows that loss of APC in excitatory forebrain neurons can disrupt the maturation and function of inhibitory GABAergic networks and, along with changes in excitatory neurons, may contribute to ISS-relevant phenotypes.

## Materials and Methods

### Animals

APC cKO mice of both sexes were generated as previously described (Mohn et al 2014). Briefly, APC was conditionally deleted by crossing male mice homozygous for the floxed APC allele (APC^fl/fl^) with female mice that are heterozygous for the floxed APC allele (APC^fl/+^) and express Cre-recombinase under the control of the CaMKIIα promoter (CaMKIIαCre). APC^fl/fl^ mice are on a mixed FVB and C57B6J background. CamKIIα mice are on a mixed C57B6/J and 129 background. This generates homozygous APC cKO mice (APC^fl/fl^/Cre+), mice heterozygous for APC in excitatory neurons (APC^fl/+^/Cre+), and WT mice (APC^fl/fl^/Cre-, APC^fl/+^/Cre-). Only homozygous cKO mice and WT mice are used in this study. To identify parvalbumin-positive cortical interneurons (PV+INs), we also integrated GAD67-eGFP (G42) mice (CB6F1/J background), in which a subset of PV+INs are labeled with green fluorescent protein (GFP+) (Chattopadhyaya et al., 2004; Hanson et al., 2019) (JAX stock #007677), into our APC cKO breeding strategy to generate triple transgenic mice. Finally, to identify both PV+ and somatostatin-positive (SST+) INs, Lhx6-GFP mice (FVB/NTac background) were also used (Tg(Lhx6-EGFP)BP221Gsat/Mmmh, RRID:MMRRC_000246-MU) which express GFP in MGE-derived neurons. Triple transgenic mice were generated that were APC cKO or WT (as above) but also contained the Lhx6-GFP gene. Cre-negative and non-APC floxed littermates were used as controls. As above, in both G42-GFP and Lhx6-GFP experiments, all APC cKO mice were homozygous knock-outs. All procedures were approved by the Tufts University Institutional Animal Care and Use Committee. Due to the complexity of the genetic manipulations used, animals were not balanced by sex, but any available animals of the appropriate genotype were used. Experimenters were blinded to genotype until after data analysis was completed.

### Brain Slice Preparation

Somatosensory cortical brain slices (400 μm) were prepared from P9, P14, and >P60 APC cKO mice or wildtype (WT) littermates as previously described (Cantu et al., 2015; Hanson et al., 2015; Pirone et al., 2017; Hanson et al., 2019; Koenig et al., 2019). Mice were anesthetized with isoflurane, decapitated and brains were rapidly removed and placed in ice-cold cutting solution consisting of (in mM): 234 sucrose, 11 glucose, 24 NaHCO_2_, 2.5 KCl, 1.25 NaH_2_PO4, 10 MgSO_4_, and 0.5 CaCl_2_. Cutting solution was constantly oxygenated with a mixture of 95%O_2_:5%CO_2_. The brain was glued to the slicing stage of Vibratome 3000 sectioning system (Leica) and slices were cut in a coronal orientation. The slices were then incubated in 32°C oxygenated artificial cerebral spinal fluid (aCSF) consisting of (in mM): 126 NaCl, 2.5 KCl, 1.25 NaH_2_PO_4_, 1 MgSO_4_, 2 CaCl_2_, 10 glucose, 26 NaHCO_2_ for 1 hour. Slices were then allowed to cool to room temperature and used for electrophysiological experiments.

Slices were placed in the recording chamber of an Olympus BX51 microscope with continuous superfusion of oxygenated aCSF maintained at 32°C, as previously described (Cantu et al., 2015; Hanson et al., 2015; Pirone et al., 2017; Hanson et al., 2019; Koenig et al., 2019). Layer 5 pyramidal neurons and G42-GFP+ interneurons were identified using a 60X water immersion objective with epifluorescence (Olympus). Electrophysiological signals were obtained using a Multiclamp 700B amplifier (Molecular Devices), low pass filtered at 1 kHz and digitized at 20kHz. Data was recorded onto a computer (Digidata 1440A, Molecular Devices) using pClamp software (Molecular Devices). Recording pipettes were pulled from borosilicate glass (3-5 MΩ, Sutter Instruments). For voltage clamp recordings, a CsMs-based internal solution was used containing (in mM): 140 CsMs, 10 HEPES, 5 NaCl, 0.2 EGTA, 5 QX314, 1.8 MgATP, 0.3 NaGTP. To isolate spontaneous and miniature excitatory postsynaptic potentials (s/mEPSCs), cells were voltage-clamped at a holding potential of -60mV. To isolate spontaneous and miniature inhibitory postsynaptic potentials (s/mIPSCs), cells were voltage-clamped at a holding potential of 0 mV. Glutamate receptors antagonists (DNQX [20 µM] and CPP [10 µM]) were included in the aCSF for IPSC recordings. Miniature E/IPSCs were recorded in the presence of 1 µM tetrodotoxin (TTX; Abcam). For current clamp recordings, internal solution contained (in mM): 130 K-gluconate, 10 HEPES, 5 KCl, 5 EGTA, 2 NaCl, 1 MgCl_2_, and 1.8 MgATP, 0.3 NaGTP. In current clamp recordings, glutamate and GABA receptor antagonists (DNQX [20 µM], CPP [10 µM], and gabazine [10 µM]) were included in the aCSF to block synaptic activity. Current injection steps were applied ranging from -200 to 400 pA in 10 or 20 pA steps. Access resistance was monitored throughout whole cell recordings and electrophysiology data were not further analyzed if the access resistance changed by greater than 20%. For s/m E/IPSCs, 2-minute recording segments were analyzed using MiniAnalysis (Synaptosoft) and average frequency and amplitude were quantified. Cumulative probability distributions were calculated by taking 100 randomly sampled events from each recorded cell and then pooling the randomly sampled data across cells in each condition.

### Interneuron Filling and Reconstruction

To reconstruct interneuron dendrites, recording pipettes were filled with an internal recording solution that contained 0.2% biocytin. Current injections from -200 to 400 pA were injected in 10pA intervals to facilitate cell filling. Then the patch pipette was slowly withdrawn from the cell, forming an outside-out patch, allowing the cell to reseal. Sections were incubated in oxygenated aCSF for at least 15 minutes and then placed in 4% paraformaldehyde (PFA) for 24 hours. After PFA fixation, sections were washed 4x in PBS and then incubated overnight in 2% BSA, 10% goat serum, 0.1% tritonX 100, and 1:50 streptavidin Cy3 (Invitrogen). The next day, sections were washed 4x in PBS, followed by 50% glycerol:PBS and 90% glycerol:PBS, for 30 min each. Slices were mounted with Fluoromount-G (Southern Biotechnology). Filled cells were imaged on an A1R confocal microscope (Nikon) with a 20x objective (Nikon). Z-stacks were taken of the cells, in 1 µm increments, to include all visible cellular processes. Three-dimensional reconstructions of dendrites and Sholl analyses were done using Imaris (Bitplane).

### G42-GFP, Lhx6-GFP, and Cl-Caspase 3 Cell Counts

Animals (P3, P5, P7, P9, P14 and >P60) were transcardially perfused with PBS followed by 4% paraformaldehyde (PFA) in 0.4M phosphate buffer. Brains were dissected and post-fixed in PFA for 24 hours, then cryopreserved in 30% sucrose solution for a minimum of 3 days. Cortical brain sections (40 μm) were collected using a cryostat (Microm, Thermo Scientific) and free-floating sections were blocked in 10% goat serum (GS), 5% bovine serum albumin (BSA), and 0.2% TWEEN20-PBS (PBST). Sections were immunolabeled with primary antibodies against GFP (1:1,000; Abcam ab13970), parvalbumin (1:2,500; Swant McAB 235), somatostatin (1:100; Millipore MAB354), and cleaved caspase 3 (1:500; Cell Signaling Technology Asp175 D3E9) in 5% GS, 1% BSA, and 0.2% PBST. All secondary antibodies were used at a ratio of 1:500 in 5% GS, 1% BSA, and 0.2% PBST (anti-chicken-GFP, Abcam ab6873; anti-mouse-Cy3, Jackson ImmunoResearch 115-165-003; anti-rat-Alexa-fluor-647, Jackson ImmunoResearch 112-605- 167; anti-rabbit-Alexa-fluor-594, Molecular Probes). Images were collected for cell counting using a Keyence epifluorescence microscope from a single plane of focus per section. Composite images of whole cortex were made using 10X objective. Illumination power and exposure were set to optimize cell body visualization and kept consistent across age and genotype. Using this imaging modality, all cells in the plane of focus were quantified. Regions of interest (ROIs) were drawn using Image J software (NIH) in primary somatosensory cortex, primary motor cortex, and infralimbic and prelimbic areas combined for prefrontal cortex, and cell numbers were counted within the ROI. The somatosensory, motor, and prefrontal (infralimbic and prelimbic areas) cortex was identified using the Allen Mouse Brain Atlas (https://mouse.brain-map.org/static/atlas). Cells within the somatosensory, motor, and prefrontal cortex were counted from the pial surface to the white matter underlying the cortex. To define the medial/lateral boundary of the ROI, we again consulted the Allen Mouse Brain Atlas to define a rectangular region safely within the anatomical borders of each region. For somatosensory cortex, this region was directly above the lateral portion of the hippocampus, for motor cortex, this region was directly above the medial portion of the hippocampus, for the prefrontal cortex, the medial cortex before the corpus callosum crosses for the prefrontal cortex was used. The ROI size was approximately 500-600K µm^2^ for all regions. There was no difference in the size of the ROI used to quantify cell density in WT as compared to cKO. A slightly smaller ROI size was used at P9, as compared to adult, consistent with a decreased brain size at that age. Cells were identified based on epifluorescence and cellular morphology of the cell body. A range of 2-5 sections, with one ROI per section, were used per animal. All the resulting densities from these ROIs were used in linear mixed-model statistical analysis, while the average of all densities from one animal were used in t-test statistical analysis. GFP+, PV+, and SST+ cell densities were calculated by dividing the number of cells by the area of the quantified ROI. Laminar density ratios were calculated by dividing superficial cortical layer densities (layers 2/3) by deep cortical layer densities (layers 4, 5/6). Cl-caspase 3 ratios were calculated by dividing the number of G42-GFP/Cl-casp+ cells by total G42-GFP+ cells in the ROI. Putative PV+IN density in the Lhx6-GFP background (Lhx6-GFP+/SST-) was calculated by subtracting Lhx6-GFP+/SST+ IN density from total Lhx6-GFP+ density.

### Statistical Analysis

At least 3 animals of each genotype and age were used per experiment. N values for each experiment are included in the figure legend. The normality of all data was tested using Shapiro-Wilk test. Student’s two sample t-test was used to compare WT and cKO mice in normally distributed data. Wilcoxon rank-sum test was used in non-normally distributed data. Both sexes were included in all experiments. At P9 and P14 animal sex was not recorded. For P60 immunolabeling and IPSC recording sex was evaluated as a biological variable for all data collected. Only mIPSC amplitude in APC cKO mice (described in results section) showed an effect of sex as a biological variable. Because most measures did not show an effect of sex as a biological variable, the challenges of sex-balancing studies using triple-transgenic mice, and previous data showing no difference in spasm and seizure phenotypes between males and females (Mohn et al., 2014a; Pirone et al., 2017), we pooled male and female data at all timepoints. This caveat is noted in the discussion. Outliers were not excluded in any datasets. For all experiments, except synaptic recording cumulative distribution analysis, α = 0.05. Averages of multiple measurements from the same animal were used in t-tests and rank-sum tests. To analyze the cumulative distribution plots of synaptic recordings, MATLAB was used to randomly sample 100 inter-event intervals (IEIs) and amplitudes per cell recording to generate distributions. A Kolmogorov-Smirnov (K-S) test was used to compare the distribution between genotypes. Alpha for K-S tests was set to 0.00001 based on a 5% false positive rate when re-sampling the same control datasets. For all experiments, we also performed linear mixed-effects modeling (LMM) on the same dataset to determine statistical significance while accounting for sources of both intra- and inter-animal variability (Aarts et al., 2014; Boisgontier and Cheval, 2016; Lau et al., 2017; Hanson et al., 2019; Koenig et al., 2019; Yu et al., 2021). LMM utilizes all measurements taken from the same animal while accounting for both intra- and inter-animal variability through the use of fixed and random effects. Fixed effects in our models included terms for genotype and interaction of genotype with another measure (i.e., current injection amplitude). Random effects in our models included terms to capture the dependency in data from the same cell/tissue and the same mouse. Example code used in RStudio (Ver. 1.2.5033) for running LMM using lme4, lmeRtest, matrix and stats packages on P9 current injections was: >currentinjectionP9 = lmer (APs ∼ current_step*genotype + (1|mouse) + (1|mouse:cell), data = currentinjectionP9_LMM, na.action = "na.omit", REML=FALSE) In which a data set (currentinjectionP9_LMM) was used to analyze the effect of genotype and current step (current_step*genotype) on action potential number (APs), and in which the inter- (1|mouse) and intra-animal (1|mouse:cell) variability was considered. For LMM, t values > 1.96 and < −1.96 we considered to be statistically significant.

## Results

### Immature PV+ IN Cell Density is Altered in APC cKO Mice using G42 GAD67-eGFP mice.

To investigate IN numbers in the developing APC cKO somatosensory cortex, we quantified the density of PV+ and somatostatin-positive (SST+) INs, medial ganglionic eminence (MGE)-derived inhibitory cells that together make up 60% of cortical GABAergic INs. Because PV protein is not expressed until approximately P12-P14 in the cerebral cortex (Solbach and Celio, 1991; de Lecea et al., 1995), we used two different genetic approaches to identify immature PV+INs, GAD67-GFP and Lhx6-GFP reporter lines. APC cKO mice were crossed with G42 mice (GAD67-eGFP), which have a GFP expression cassette inserted under the control of the GAD67 promoter (Chattopadhyaya et al., 2004). Multiple studies confirm that a subset of PV+INs express GFP as early as P3 in G42 mice, well before the expression of PV protein (Chattopadhyaya et al., 2004; Southwell et al., 2012; Hanson et al., 2019). G42-GFP+, PV+, and SST+ cells were quantified in the somatosensory cortex at P9 and in adult (>P60) mice. G42-GFP+ IN cell density was significantly lower at P9 in the cKO mice (1.61 ± 0.22 cells/10,000 µm^2^) compared to WT littermates (2.64 ± 0.18 cells/10,000 µm^2^; Fig. 1b,c). No immunohistochemical labeling of PV+INs was seen in either genotype at P9, consistent with lack of PV expression at this age (data not shown). SST+ IN density was similar between P9 APC cKO (2.39 ±0.34 cells/10,000 µm2) and WT (2.75 ±0.57 cells/10,000 µm^2^) mice (Fig. 2a,b).

**Fig 1.**
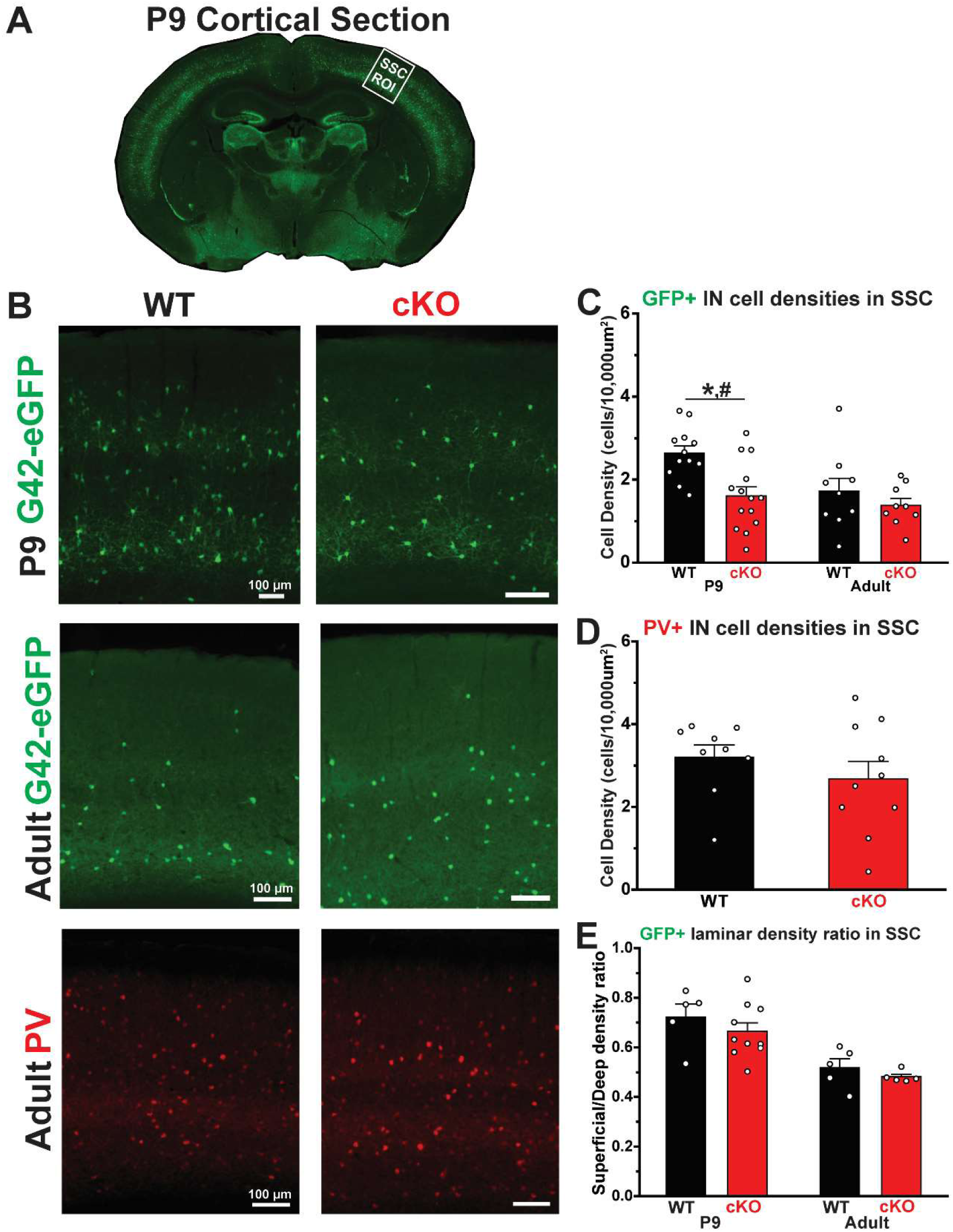
`Immature PV+IN density, as measured using G42-GFP mice, is reduced in the somatosensory cortex of APC cKO mice at P9`. `A.` Example ROI drawn in primary somatosensory cortex (SSC) of a P9 mouse brain section. `B.` Representative fluorescent images in SSC of G42-GFP+ (green, labeling a subset of PV+INs) cells at (top) P9 and (middle) adult (>P60) and (bottom) PV+ immunolabeled (red) cells in adult for WT and cKO mice. Scale bar = 100µm. `C.` Bar graph of average G42-GFP+ IN cell density per animal at P9 for WT (black bar, n=12 animals) and cKO (red bar, n=14 animals) mice (p=1.60e-3, t(24)=3.56, two sample t-test; p=9.86e-4, t(61)=3.70, LMM) and adult WT (n=9 animals) and cKO (n=9 animals; p=0.36, t(16)=0.95, two sample t-test; p=0.27, t(39)=1.13, LMM). `D.` Bar graph of average PV+IN densities per animal in adult for WT (black bar, n=9 animals) and cKO (red bar, n=10 animals) mice (p=0.33, t(17)=1.00, two sample t-test; p=0.92, t(23)=0.11, LMM). `E.` Bar graph of the laminar density ratio, which represents the ratio of superficial layer densities to deep layer densities, of G42-GFP+ INs at P9 in WT (black bar, n=5 animals) and cKO (red bar, n=10 animals) mice (p=0.35, t(13)=0.96, two sample t-test; p=0.21, t(39)=1.32, LMM) and adult WT (n=5 animals) and cKO (n=5 animals; p=0.36, t(8)=0.98, two sample t-test; p=033, t(19)=1.01, LMM). *p<0.05, two-sample t test. #p<0.05, LMM, effect of genotype. Each data point represents the average ratio for one animal. Error bars indicate SEM.

**Fig 2.**
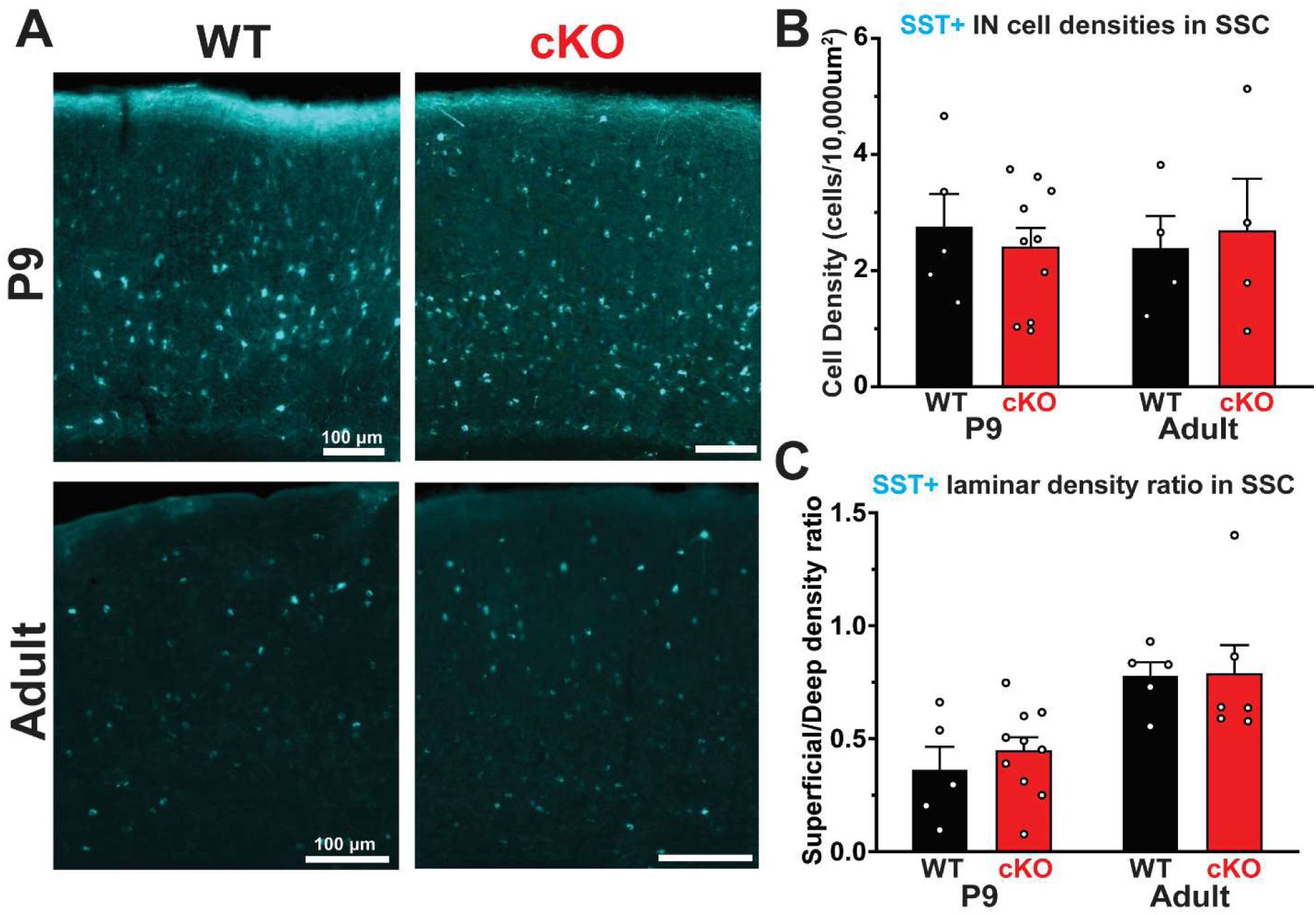
`SST+ IN density does not differ in the somatosensory cortex of APC cKO mice`. `A.` Representative fluorescent images in SSC of SST+ immunolabeled (cyan) cells at (top) P9 and (bottom) adult in WT and cKO mice. Scale bar = 100µm. `B.` Bar graph of average SST+ IN cell density per animal at P9 for WT (black bar, n=5 animals) and cKO (red bar, n=10 animals) mice (p=0.58, t(13)=0.57, two sample t-test; p=0.56, t(36)=0.60, LMM) and adult WT (n=4 animals) and cKO (n=4 animals; p=0.79, t(6)=0.28, two sample t-test; p=0.74, t(8)=-0.34, LMM). `C.` Bar graph of the laminar density ratio, which represents the ratio of superficial layer densities to deep layer densities, of SST+ INs at P9 in WT (black bar, n=5 animals) and cKO (red bar, n=10 animals) mice (p=0.47, t(13)=-0.74, two sample t-test; p=0.41, t(39)=-0.84, LMM) and adult WT (n=5 animals) and cKO (n=6 animals; p=0.95, t(9)=-0.06, two sample t-test; p=0.94, t(20)=-0.08, LMM). *p<0.05, two-sample t test. #p<0.05, LMM, effect of genotype. Each data point represents the average ratio for one animal. Error bars indicate SEM.

In adult mice, there was no significant difference in the density of G42-GFP+ cells (cKO density 1.38 ± 0.17 cells/10,000 µm^2^; WT density 1.72 ± 0.32 cells/10,000 µm^2^; Fig. 1b,c), PV+ immunolabeled INs (cKO density 2.68 ± 0.42 cells/10,000 µm^2^; WT density 3.20 ± 0.30 cells/10,000 µm^2^; Fig. 1b,d), or SST+ INs (cKO density 2.67 ± 0.90 cells/10,000 µm^2^; WT density 2.37 ±0.57 cells/10,000 µm^2^; Fig. 2a,b) between APC cKO and WT mice. The laminar density between superficial (layers 2/3) and deep (layers 4, 5/6) cortical layers did not differ for G42-GFP+ interneurons at P9 (cKO ratio 0.66 ± 0.03; WT ratio 0.72 ± 0.05) or adult (cKO ratio 0.48 ± 0.01; WT ratio 0.52 ± 0.04; Fig. 1e). The laminar distribution of SST+ INs was also not altered (P9: cKO ratio 0.44 ± 0.06; WT ratio 0.36 ± 0.11; Adult: cKO ratio 0.78 ± 0.13; WT ratio 0.77 ± 0.06; Fig. 2c). To determine whether similar reductions in G42-GFP+ INs were seen in other regions, we also examined motor (MC) and prefrontal cortex (PFC) in P9 and adult mice (Fig. 3). In APC cKO mice, G42-GFP+ IN density was decreased at P9, but not adults in the MC (P9: WT 1.95 ± 0.15 cells/10,000 µm^2^, cKO 1.14 ± 0.14 cells/10,000 µm^2^; Adult: WT 1.62 ± 0.27 cells/10,000 µm^2^, cKO 1.13 ± 0.17 cells/10,000 µm^2^, Fig. 3a) and PFC (P9: WT 1.26 ± 0.14 cells/10,000 µm^2^, cKO 0.77 ± 0.12 cells/10,000 µm^2^; Adult: WT 1.13 ± 0.26 cells/10,000 µm^2^, cKO 0.61 ± 0.09 cells/10,000 µm^2^, Fig. 3d), similar to what was we report in SSC. At P9, SST+ IN density was similar in WT and APC cKO mice in both MC (WT 3.42 ± 0.67 cells/10,000 µm^2^, cKO 2.81 ± 0.36 cells/10,000 µm^2^, Fig. 3c) and PFC (WT 1.67 ± 0.32 cells/10,000 µm^2^, cKO 2.52 ± 0.79 cells/10,000 µm^2^, Fig. 3f) PV+IN and SST+ IN density was not altered in MC in adults (PV: WT 2.72 ± 0.30 cells/10,000 µm^2^, cKO 2.36 ± 0.47 cells/10,000 µm^2^; SST: WT 2.24 ± 0.66 cells/10,000 µm^2^, cKO 2.60 ± 0.92 cells/10,000 µm^2^, Fig. 3b,c), similar to the SSC, but both were reduced in adult PFC (PV: WT 2.78 ± 0.48 cells/10,000 µm^2^, cKO 1.23 ± 0.24 cells/10,000 µm^2^; SST: WT 2.28 ± 0.36 cells/10,000 µm^2^, cKO 1.21 ± 0.17 cells/10,000 µm^2^, Fig. 3e,f), consistent with a previous study (Pirone et al., 2018).

**Figure 3.**
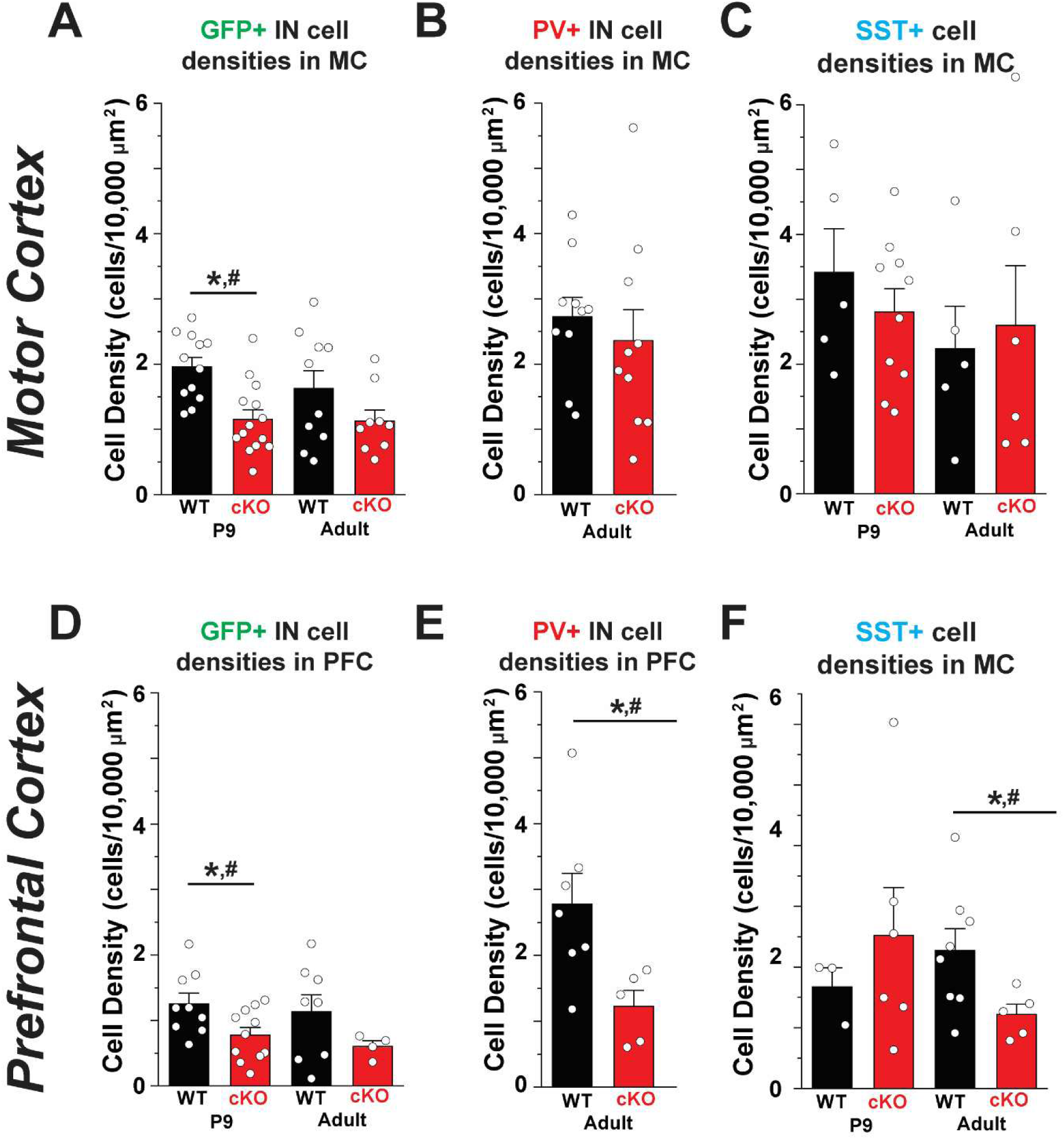
`Changes in G42-GFP+, PV+, and SST+ INs in the motor and prefrontal cortex. A.` Bar graph of the average G42-GFP+ IN density in the motor cortex in P9 WT (n=12 animals) and cKO (n=14 animals; p=6.92e-4, t(24)=3.89, two sample t-test; p=6.24e-4, t(73)=3.86, LMM); and adult WT (n=10 animals) and cKO (n=9 animals; p=0.15, t(17)=1.52, two sample t-test; p=0.14, t(40)=1.55, LMM). `B.` PV+ immunolabeled IN densities in adult WT motor cortex (n=10 animals) and cKO (n=10 animals; p=0.53, t(18)=0.65, two sample t-test; p=0.76, t(36)= -0.32, LMM) mice in the motor cortex. `C.` Bar graph of the average SST+ IN density in P9 WT motor cortex (n=5 animals) and cKO (n=10 animals; p=0.39, t(13)=0.89, two sample t-test; p=0.40, t(60)=0.86, LMM); and adult WT (n=5 animals) and cKO (n=6 animals; p=0.77, t(9)=0.30, two sample t-test; p=0.69, t(22)= -0.41, LMM) mice in the motor cortex. `D.` Bar graph of the average G42-GFP+ IN density in P9 WT (n=9 animals) and cKO (n=11 animals; p=0.02, t(18)=2.45, two sample t-test; p=0.02, t(21)=2.66, LMM); and adult WT (n=8 animals) and cKO (n=4 animals; p=0.19, t(10)=1.40, two sample t-test; p=0.15, t(18)=1.52, LMM) prefrontal cortex. `E.` PV+ immunolabeled IN densities in adult WT(n=7 animals) and cKO (n=5 animals; p=0.03, t(10)=2.60, two sample t- test; p=0.01, t(13)=3.02, LMM) mice in the prefrontal cortex. `F.` Bar graph of the average SST+ IN densities in P9 WT (n=3 animals) and cKO (n=6 animals; p=0.49, t(7)=0.72, two sample t-test; p=0.47, t(10)= -0.76, LMM); and in adult WT (n=8 animals) and cKO (n=5 animals; p=4.95e-2, t(11)=2.21, two sample t-test; p=0.03, t(25)=2.33, LMM) mice in the prefrontal cortex. *p<0.05, two-sample t test. #p<0.05, LMM, effect of genotype. Each data point represents the average ratio for one animal. Error bars indicate SEM.

### Immature PV+ IN Cell Density is Altered in APC cKO Mice using Lhx60-GFP mice.

To confirm that immature PV+IN density is lower in APC cKO mice at P9, we repeated this experiment using Lhx6-GFP mice, a different genetic reporter line in which all MGE-derived INs (PV+ and SST+) are labeled with GFP. Because SST protein is robustly expressed early in cortical development, immature PV+INs can be identified as cells that are Lhx6-GFP+ but SST immuno- negative (Liodis et al., 2007; Alroy et al., 2008). The density of all Lhx6-GFP+ IN (representing both PV+ and SST+ INs) was significantly decreased in the SSC for APC cKO mice at P9 (cKO density 3.47 ± 0.47 cells/10,000 µm^2^; WT density 5.33 ± 0.37 cells/10,000 µm^2^; Fig. 4a,b). Next, we quantified the density of Lhx6 INs that are SST+ (Lhx6+/SST+) at P9 and found there was no difference between WT and cKO (cKO density 0.56 ± 0.10 cells/10,000 µm^2^; WT density 0.68 ± 0.09 cells/10,000 µm^2^; Fig 4a,c). The density of immature PV+INs (Lxh6-GFP+/SST-), was calculated by subtracting the Lhx6-GFP+/SST+ cell density from total Lhx6-GFP+ cell density. Using this approach, we found that immature PV+IN density was again decreased in APC cKO mice at P9 (cKO density 2.92 ± 0.40 cells/10,000 µm2; WT density 4.64 ± 0.43 cells/10,000 µm2; Fig. 4a,d). In adult APC cKO mice we found that there was no difference in either Lhx6-GFP+ (cKO density 2.51 ± 0.27 cells/10,000 µm2; WT density 3.46 ± 0.41 cells/10,000 µm2; Fig. 4e,f) or in double positive PV+/Lhx6-GFP+ cells (cKO density 1.06 ± 0.15 cells/10,000 µm2; WT density 1.52 ± 0.19 cells/10,000 µm2; Fig. 4e,g; please note: this results in labeling of a subset of PV+IN that are GFP+ in the Lhx6-GFP line). Together, these studies show that immature PV+INs are decreased in the P9 somatosensory cortex in APC cKO mice. For the subsequent experiments, G42 reporter mice were used to identify immature PV+INs.

**Fig 4.**
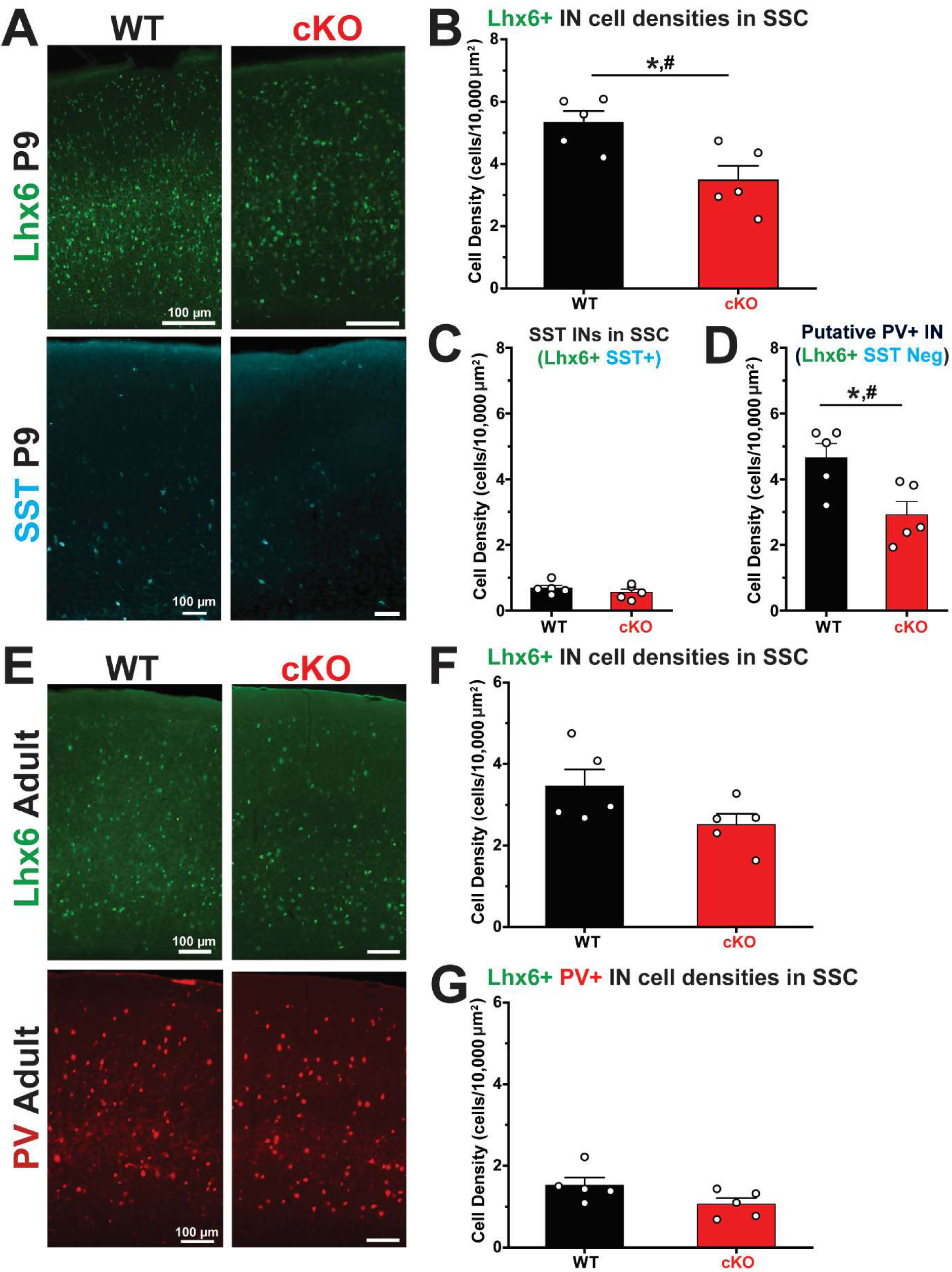
`Immature PV+IN density, as measured using Lhx6-GFP mice, is reduced in the somatosensory cortex of APC cKO mice at P9`. `A.` Representative fluorescent images in SSC of Lhx6+ cells (top, green, labeling MGE-derived INs) and SST+ cells (bottom, cyan) at P9 in WT and cKO mice. Scale bar = 100µm scale. `B.` Bar graph of average Lhx6+ IN (all MGE-derived INs) density per animal from SSC at P9 in WT (black, n=5 animals) and cKO (red, n=5 animals; p=0.01, t(8)= 3.12, two sample t-test; p=6.59e-3, t(15)=3.41, LMM) mice. `C.` Bar graph of average Lhx6+/SST+ IN (genetically Lhx6+ and SST immune-positive) density per animal at P9 in WT (n=5 animals) and cKO (n=5 animals; p=0.35, t(8)=0.99, two sample t-test; p=0.29, t(15)=1.10, LMM) mice. `D.` Bar graph of average putative PV+IN (genetically Lhx6+ and SST immune- negative) density per animal at P9 in WT (n=5 animals) and cKO (n=5 animals; p=0.02, t(8)= 2.92, two sample t-test; p=0.01, t(15)=3.19, LMM) mice. `E.` Representative fluorescent images in SSC of Lhx6+ cells (top) and PV+ immunolabeled cells (red, bottom) in adult WT and cKO mice. `F.` Bar graph of average Lhx6+ IN density per animal from SSC in adult WT (n=5 animals) and cKO (n=5 animals; p=0.09, t(8)= -1.94, two sample t-test; p=0.06, t(14)=2.12, LMM) mice. `G.` Bar graph of average Lhx6+/PV+IN density per animal from immunolabeling in adult WT (n=5 animals) and cKO (n=5 animals; p=0.09, t(8)= -1.92, two sample t-test; p=0.06, t(14)=2.10, LMM) mice. *p<0.05, two-sample t test. #p<0.05, LMM, effect of genotype. Each data point represents the average density for one animal. Error bars indicate SEM.

### Immature G42-GFP+ PV+ INs Undergo Disproportionate Developmental Apoptosis in APC cKO Mice

Cortical INs are produced in excess during embryonic development (Denaxa et al., 2018) and their final numbers are established through a process of intrinsic apoptosis wherein death occurs approximately 12 days after genesis, unless prevented by external neuronal activity (Southwell et al., 2012; Wong et al., 2018). Although the threshold of excitatory input necessary to confer survival is not established, IN survival is improved by calcium influx and action potential generation (Priya et al., 2018). MGE-derived interneuron cell death peaks at approximately P7-P9, the time of peak behavioral spasms in APC cKO mice. We hypothesized that decreased PV+IN density in neonatal APC cKO cortex may arise from altered developmental apoptosis.

To investigate the developmental apoptosis of PV+INs during early development, we used an antibody against cleaved caspase 3 (Cl-CASP3+), a marker of apoptosis. We examined 3 developmental time windows: P3-P5 (before behavioral spasm onset), P7-P9 (during the peak of behavioral spasms), and P14 (after spasms stop) (Pirone et al., 2017). G42-GFP+ IN density was decreased in P7-P9 APC cKO mice (cKO density 1.51 ± 0.28 cells/10,000 µm^2^; WT density 2.30 ± 0.23 cells/10,000 µm^2^; Fig. 5b), consistent with data in Figures 1. The density of G42-GFP+ INs that co-labeled for Cl-CASP3+ (G42-GFP+/Cl-CASP3+) was unchanged at from P3-5 and P7-9, but was significantly decreased at P14 (cKO density 0.44 ± 0.09 cells/10,000 µm^2^; WT density 1.05 ± 0.26 cells/10,000 µm^2^; Fig. 5c). Because the number of G42-GFP+ INs was not equivalent in WT and APC cKO mice, we next asked if the proportion of G42-GFP+ cells undergoing apoptosis was altered in APC cKOs. At P3-P5, the percentage of G42-GFP+ cells that were co- labeled by Cl-CASP3+ was similar in APC cKO and WT mice (cKO ratio 0.31 ± 0.05; WT ratio 0.27 ± 0.04; Fig. 5d). However, from P7-P9 there were significantly more G42-GFP+ INs that were Cl-CASP3+ in APC cKO mice, as compared to WT (cKO ratio 0.71 ± 0.05; WT ratio 0.50 ± 0.05; Fig. 5a,d). This increase is likely driven by the fact that there are less G42-GFP+ INs in the SSC at this time point (Fig. 5b,c). At P14, APC cKO mice had significantly fewer G42-GFP+ cells that were Cl-CASP3+ (cKO ratio 0.28 ± 0.05; WT ratio 0.55 ± 0.08; Fig. 4d), suggesting that developmental cell death in PV+INs ends earlier in the cKO. Because developmental apoptosis of PV+INs is activity dependent, we next asked whether excitation of PV+INs was disrupted in early development in APC cKO mice.

**Fig 5.**
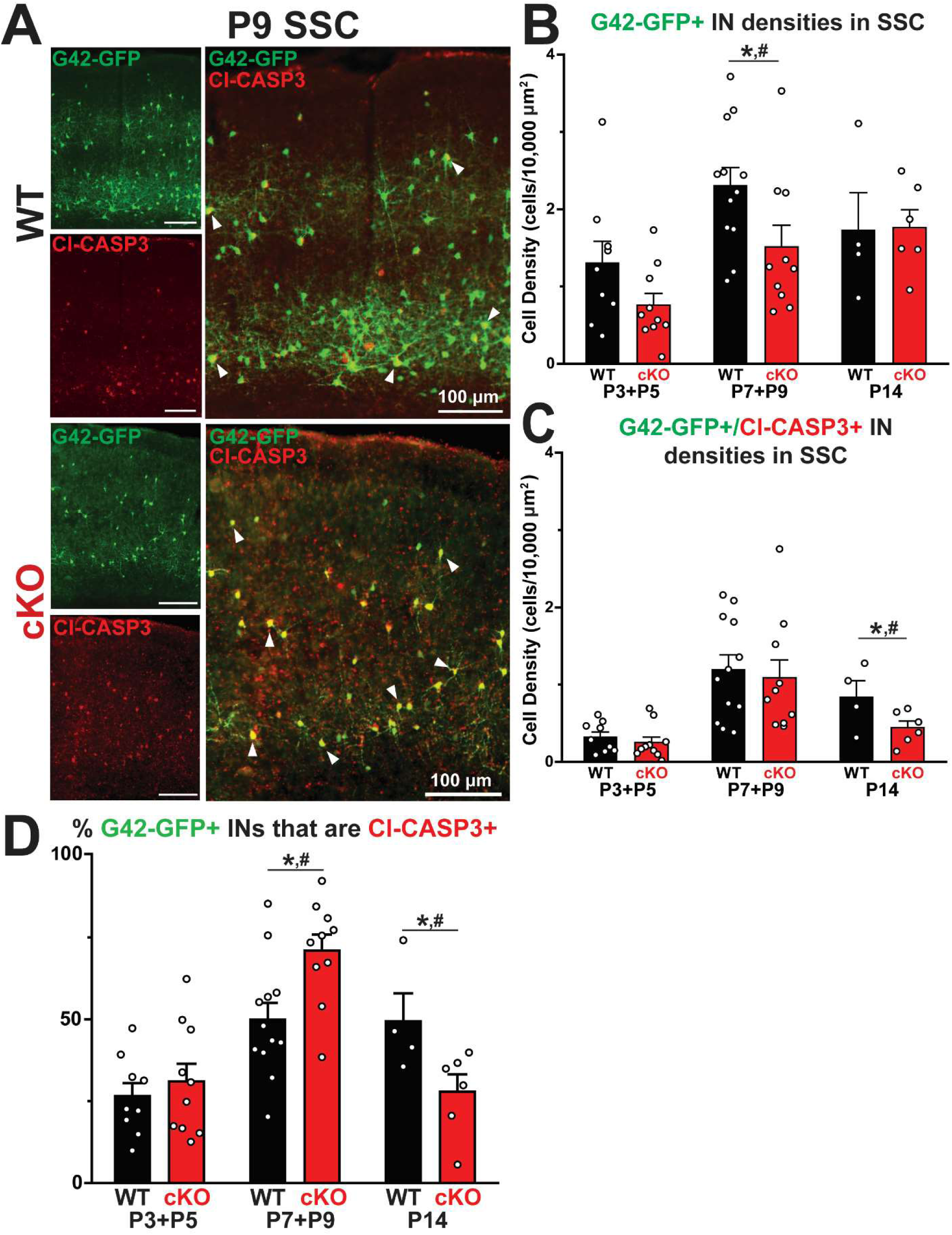
`APC cKO mice show disrupted developmental IN apoptosis in G42 labeled cells. A.` Representative fluorescent images in SSC of G42-GFP+ (green) cells, Cl-Caspase3 (Cl- CASP+, red) immunolabeled cells, and overlays from P9 (top) WT and (bottom) cKO mice. Scale bar = 100µm. Yellow cells indicate a G42-GFP+ IN co-labeled with Cl-CASP antibody (right) and white arrows indicate representative colabeled cells. `B.` Bar graph of the average G42-GFP+ IN density in P3-P5 WT (n=9 animals) and cKO (n=10 animals; p=0.10, t(17)= -1.75, two sample t- test; p=0.07, t(65)=1.94, LMM), P7-P9 WT (n=12 animals) and cKO (n=10 animals; p=0.04, t(20)=2.19, two sample t-test; p=0.04, t(40)=2.18, LMM), and P14 WT (n=5 animals) and cKO (n=6 animals; p=0.79, t(9)= -0.27, two sample t-test; p=0.77, t(15)=0.30, LMM) mice. `C.` Bar graph of the average G42-GFP+, Cl-CASP3+ IN density in P3-P5 WT (n=9 animals) and cKO (n=10 animals; p=0.47, t(17)= -0.74, two sample t-test; p=0.10, t(65)=1.74, LMM), P7-P9 WT (n=12 animals) and cKO (n=10 animals; p=0.72, t(20)=0.37, two sample t-test; p=0.85, t(40)=0.20, LMM), and P14 WT(n=5 animals) and cKO (n=6 animals; p=0.04, t(9)= -2.35, two sample t-test; p=0.02, t(15)=2.59, LMM) mice. `D.` Bar graph of the average percentage G42-GFP+ cells that are Cl-CASP+ in P3-P5 WT (black bar, n=9 animals) and cKO (red bar, n=10 animals; p=0.52, t(17)=0.66, two sample t-test; p=0.94, t(59)=-0.08, LMM), P7-P9 WT (n=12 animals) and cKO (n=10 animals; p=8.62e-3, t(20)= -2.91, two sample t-test; p=2.53e-3, t(40)=-3.43, LMM), and P14 WT (n=5 animals) and cKO (n= 6 animals; p=0.02, t(9)= -2.79, two sample t-test; p=0.01, t(16)=3.16, LMM). *p<0.05, two-sample t test. #p<0.05, LMM, effect of genotype. Each data point represents the average density or ratio for one animal. Error bars indicate SEM.

### Synaptic Excitation of PV+ Interneurons is Increased in APC cKO Mice

Whole cell voltage clamp recordings were made from layer 5 G42-GFP+ INs in APC WT and cKO mice at P9, P14, the ages of altered developmental apoptosis, and in adults (>P60). Spontaneous excitatory post-synaptic currents (sEPSCs) were recorded, and frequency and amplitude were quantified. At all ages, the mean frequency of sEPSCs onto PV+INs was increased in APC cKOs compared to WT mice (P9 cKO 2.70 ± 0.55 Hz; P9 WT 0.97 ± 0.12 Hz; P14 cKO 8.73 ± 1.31 Hz; P14 WT 4.96 ± 0.98 Hz; P60 cKO 6.46 ± 1.87 Hz; P60 WT 2.65 ± 0.41; Fig. 6). In addition, the cumulative distribution of sEPSC inter-event intervals was significantly shifted to the left, consistent with more frequent synaptic excitation (Fig. 6d,e,f). Mean sEPSC amplitude was similar in APC cKOs and WT mice at all ages (P9 cKO 18.5 ± 1.59 pA; P9 WT 22.8 ± 2.88 pA; P14 cKO 16.1 ± 1.94 pA; P14 WT 19.7 ± 2.47 pA; P60 cKO 23.8 ± 4.49 pA; P60 WT 20.6 ± 2.80 pA; Fig. 6), but the cumulative distribution of sEPSC amplitudes was left-shifted at P9 and P14, suggesting smaller sEPSCs in APC cKO mice (Fig. 6d,e).

**Fig 6.**
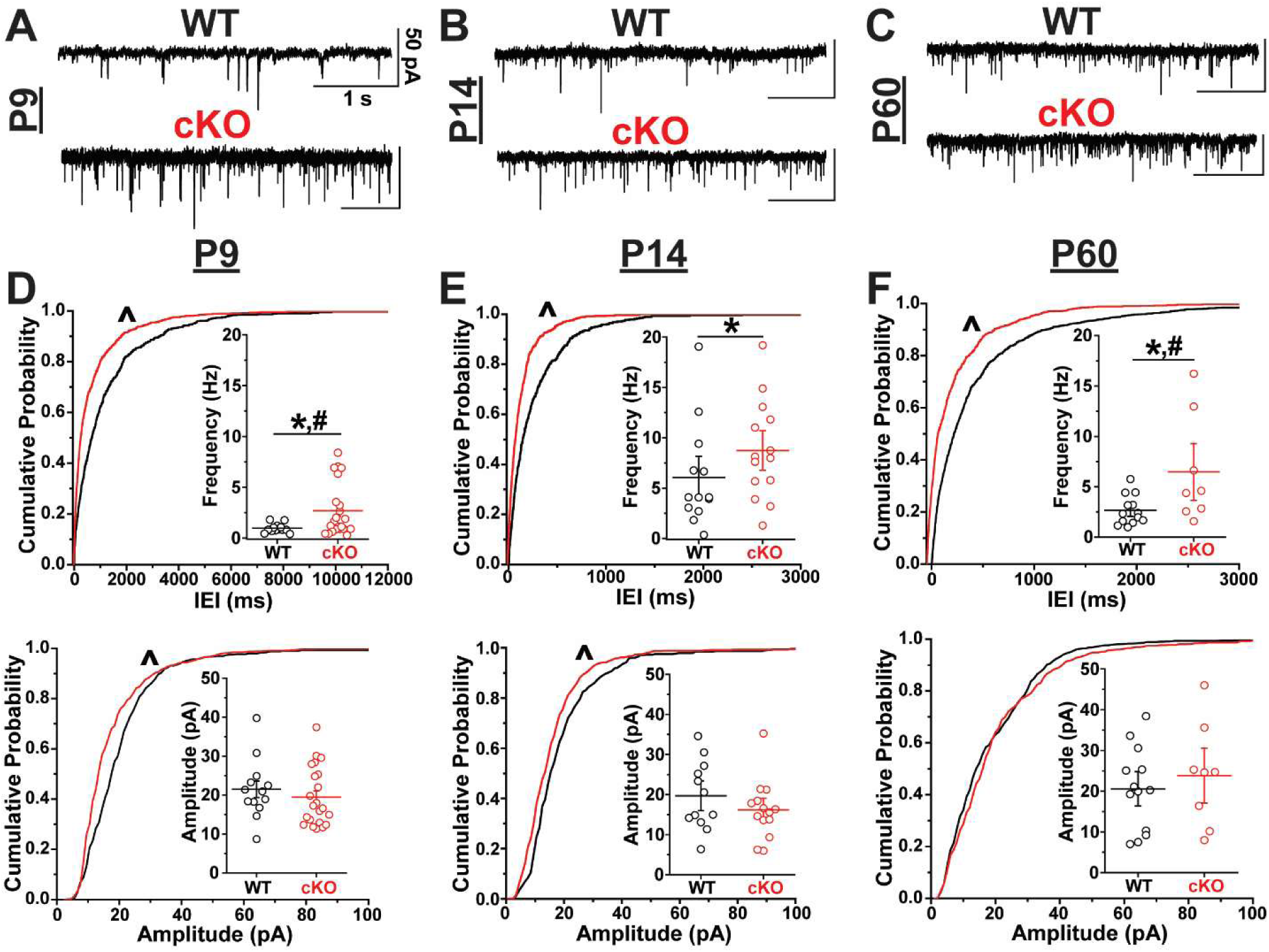
`Increased sEPSCs onto G42-GFP+ INs in the cortex of the APC cKO.` Example spontaneous excitatory post-synaptic current (sEPSC) traces from voltage-clamped, L5 G42- GFP+ INs held at -60 mV at `A.` P9, `B.` P14, and `C.` P60 for WT (black) and cKO (red) mice. All recordings made in somatosensory cortex. Scale bars = 50 pA, 1 sec. `D.` Cumulative probability plots of inter-event intervals (p=5.23e-19, Kolmogorov-Smirnov (K-S) test) with inset of mean frequency for WT (black, n=13 cells, 4 animals) and cKO (red, n=22 cells, 6 animals; p=0.03, t(32)= -2.32, two sample t-test; p=0.04, t(28)=-2.19, LMM) and amplitude (p=5.36e-14, K-S test) with inset of mean amplitude for WT (n=13 cells, 4 animals) and cKO (n=18 cells, 6 animals; p=0.17, t(29)=1.40, two sample t-test; p=0.23, t(30)=1.22, LMM) at P9. `E.` Cumulative probability plots of inter-event intervals (p=1.85e-12, K-S test) with inset of mean frequency for WT (n=14 cells, 6 animals) and cKO (n=12 cells, 5 animals; p=0.03, t(24)= -2.25; p=0.49, t(22)=-0.73, LMM) and amplitude (p=1.63e-5; K-S test) with inset of mean amplitude for WT (n=12 cells, 6 animals) and cKO (n=14 cells, 5 animals; p=0.27, t(24)=1.14, two sample t-test; p=0.40, t(22)=0.88, LMM) at P14. `F.` Cumulative probability plots of inter-event intervals (p=6.93e-13, K-S test) with inset of mean frequency for WT (n=13 cells, 5 animals) and cKO (n=8 cells, 4 animals; p=0.02, t(19)= - 2.48, two sample t-test; p=0.02, t(16)=-2.57, LMM) and amplitude (p=0.02, K-S test) with inset of mean amplitude for WT (n=13 cells, 5 animals) and cKO (n=8 cells, 4 animals; p=0.52, t(19)=- 0.65, two sample t-test; p=0.95, t(16)=0.06, LMM) at P60. ^p<0.00001, KS test. *p<0.05, two-sample t test. #p<0.05, LMM, effect of genotype. Each data point represents one cell. Data for inset graphs are mean ± SEM.

We next quantified action potential-independent miniature excitatory post-synaptic currents (mEPSCs) in layer 5 G42-GFP+ INs. At P9, the mean frequency of mEPSCs (cKO 2.60 ± 0.44 Hz; WT 0.89 ± 0.20 Hz; Fig. 7a,d) was significantly increased and the cumulative distribution of mEPSC inter-event intervals was significantly left-shifted in APC cKO mice (Fig. 7d). At P14 and P60, however, mEPSCs frequency was equivalent in APC cKO and WT mice (P14 cKO 5.74 ± 1.19 Hz; P14 WT 5.73 ± 1.01 Hz; Fig. 7b,e; P60 cKO 4.46 ± 1.14 Hz; P60 WT 3.73 ± 0.61 Hz; Fig. 7c,f). mEPSC amplitude was equivalent in APC cKOs and WTs at all ages as well (P9 cKO16.3 ± 1.36 pA; P9 WT 15.7 ± 1.30 pA; P14 cKO 16.1 ± 1.13 pA; P14 WT 16.3 ± 1.20 pA; P60 cKO 12.2 ± 1.44 pA; P60 WT 12.8 ± 1.06 pA; Fig. 7). In conclusion, G42-GFP+ INs in APC cKOs receive an increased frequency of sEPSCs at all ages examined, but mEPSCs frequency onto INs was increased only at P9. This is consistent with known increases synaptic excitation in APC cKOs (Pirone et al., 2017).

**Fig 7.**
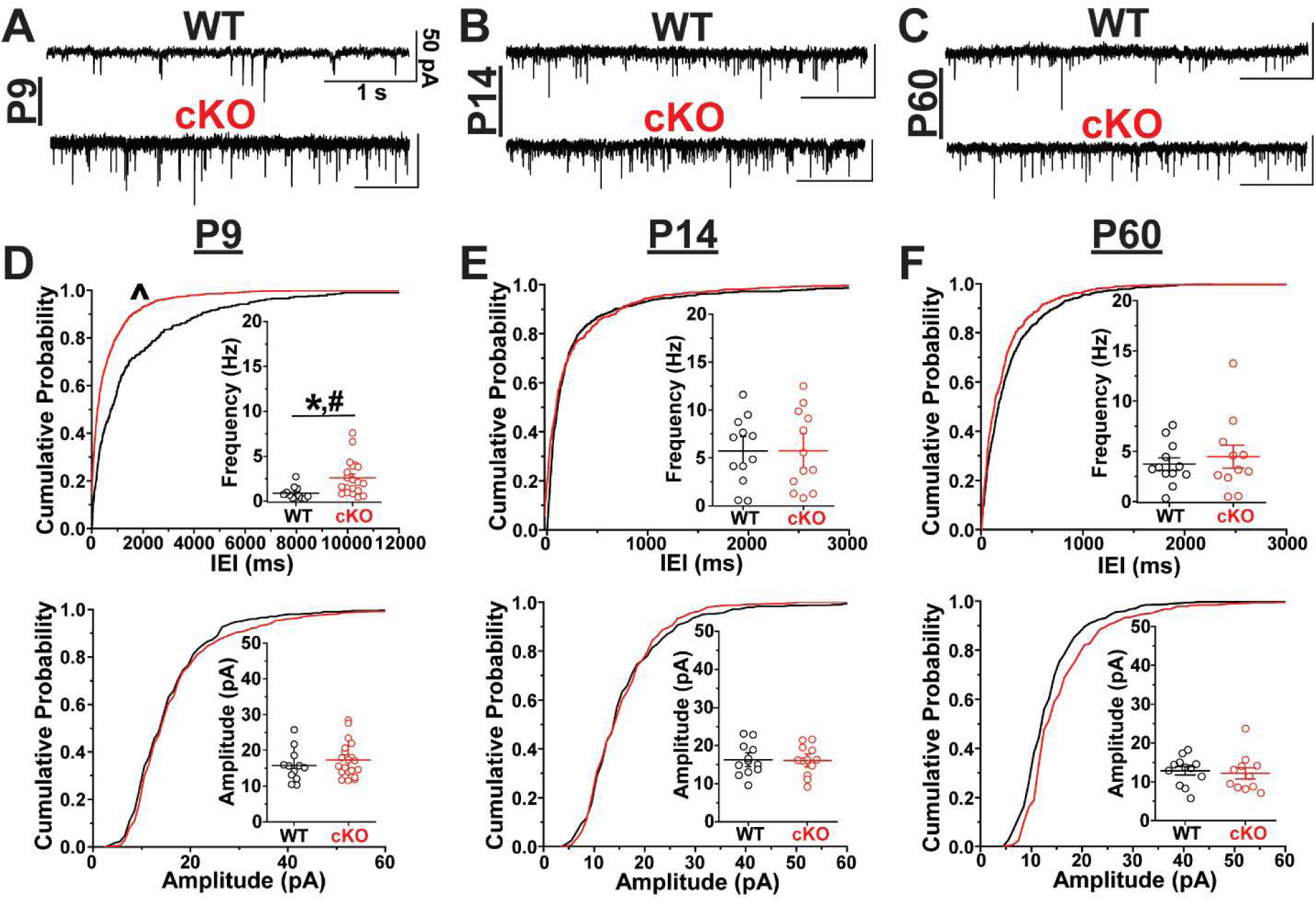
`Increased mEPSCs onto G42-GFP+ INs at P9 in the APC cKO.` Example miniature excitatory post-synaptic current (mEPSC) traces from voltage-clamped, L5 G42-GFP+ INs held at -60 mV at `A.` P9, `B.` P14, and `C.` P60 for WT (black) and cKO (red) mice. All recordings made in somatosensory cortex. Scale bars = 50 pA, 1 sec. `D.` Cumulative probability plots of inter-event intervals (p=9.67e-16, K-S test) with inset of mean frequency for WT (black, n=12 cells, 3 animals) and cKO (red, n=20 cells, 5 animals; p=0.01, t(24)= -2.77, two sample t-test; p=0.02, t(27)=-2.71, LMM) and amplitude (p=0.03, K-S test) with inset of mean amplitude for WT (n=12 cells, 3 animals) and cKO (n=13 cells, 5 animals; p=0.75, t(23)=-0.33, two sample t-test; p=0.10, t(27)=- 1.70, LMM) at P9. `E.` Cumulative probability plots of inter-event intervals (p=0.10, K-S test) with inset of mean frequency for WT (n=12 cells, 6 animals) and cKO (n=12 cells, 6 animals; p=0.99, t(22)=-0.01, two sample t-test; p=0.54, t(21)=0.64, LMM) and amplitude (p=0.76, K-S test) with inset of mean amplitude for WT (n=12 cells, 6 animals) and cKO (n=12 cells, 6 animals; p=0.90, t(22)=0.13, two sample t-test; p=0.77, t(21)=0.30, LMM) at P14. `F.` Cumulative probability plots of inter-event intervals (p=1.20e-3, K-S test) with inset of mean frequency for WT (n=12 cells, 4 animals) and cKO (n=11 cells, 3 animals; p=0.57, t(21)=-0.58, two sample t-test; p=0.52, t(17)=- 0.67, LMM) and amplitude (p=2.20e-3, K-S test) with inset of mean amplitude for WT (n=12 cells, 4 animals) and cKO (n=11 cells, 3 animals; p=0.71, t(21)=0.37, two sample t-test; p=0.88, t(17)=0.16, LMM) at P60. ^p<0.00001, KS test. *p<0.05, two-sample t test. #p<0.05, LMM, effect of genotype. Each data point represents one cell. Data for inset graphs are mean ± SEM.

### Dendrite Morphology is Altered in Immature PV+ Interneurons in APC cKO Mice

Cortical excitation guides PV+IN morphological maturation, including dendritogenesis (Murase et al., 2002; Konur and Ghosh, 2005; Close et al., 2012; Wamsley and Fishell, 2017). Since PV+INs receive increased excitation in APC cKO mice, we suspected that their dendritic morphology may also be altered. To address this question, layer 5 G42-GFP+ INs were biocytin filled, their dendrites were morphologically reconstructed, and the number of dendritic intersections at increasing radii from the soma was quantified using Scholl analysis. To analyze P9 dendritic morphology with LMM, we fit a piecewise linear mixed effects model including a knot, or change in slope, at a radius of 60 µm (Fig. 8a,b) for data collected from 0 to 300 µm from the soma. The piecewise linear mixed effects model showed that for radii less than 60 µm, the number of intersections increase more quickly for cKO mice than for WT mice at P9 (cKO slope = 0.28 µm/intersections, WT slope = 0.20 µm/intersections) and no difference from 60-300 µm at P9 (cKO slope = -0.13 µm/intersections, WT slope = -0.10 µm/intersections). G42+ INs morphology was similar in WT and APC cKO when examined at P14 (Fig. 8c,d) and P60 (Fig. 8e,f). The increase in dendritic complexity at P9 is consistent with increased mEPSC frequency (Fig. 7a,d), as more dendritic processes may be able to receive more excitatory synaptic input.

**Fig 8.**
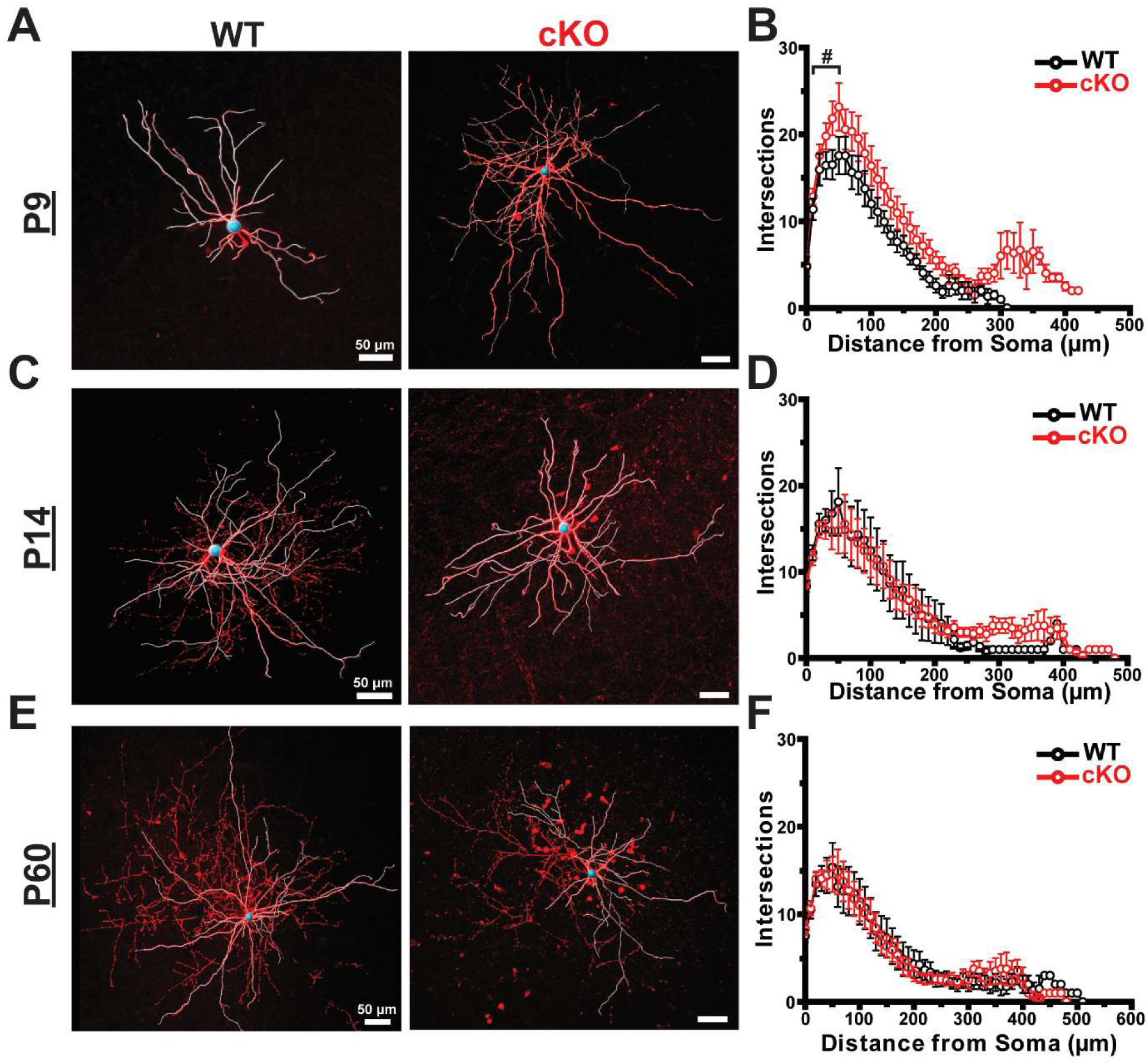
`Increased dendritic complexity of P9 G42-GFP+ INs in the APC cKO. A.` Representative confocal dendritic reconstructions of biocytin-filled G42-GFP+ INs from L5 somatosensory cortex at P9 in WT and cKO mice, 50µm scale. Blue sphere indicates the soma and white traces indicate dendrites. `B.` Scholl’s analysis plot showing the number of dendritic intersections at increasing radii from the soma for biocytin-filled G42-GFP+ INs at P9 in WT (black, n=18 cells, 5 animals) and cKO (red, n=19 cells, 4 animals) mice. For radii 0-60 µm, p=2.25e-4, t(839)=-3.71, LMM. For radii 60-300 µm, p=0.80, t(839)=0.25, LMM. `C.` Representative confocal dendritic reconstructions of biocytin-filled G42-GFP+ INs from L5 SSC at P14 in WT and cKO mice, 50µm scale. `D.` Scholl’s analysis plot showing the number of dendritic intersections at increasing radii from the soma for biocytin-filled G42-GFP+ INs at P14 in WT (n=14 cells, 5 animals) and cKO (n=12 cells, 3 animals) mice. `E.` Representative confocal dendritic reconstructions of biocytin-filled G42-GFP+ INs from L5 SSC at P60 in WT (n=12 cells, 3 animals) and cKO (n=10 cells, 4 animals) mice, 50µm scale. `F.` Scholl’s analysis plot showing the number of dendritic intersections at increasing radii from the soma for biocytin-filled G42-GFP+ INs at P60 in WT and cKO mice. #p<0.05, LMM, effect of genotype. Each data point represents the average number of intersections per genotype. Error bars indicate SEM.

### Dynamic Developmental Changes in Synaptic Inhibition in APC cKO Mice

Because the excitation of PV+INs is enhanced in APC cKOs, we suspected that inhibitory output may also be altered. We therefore made whole-cell recordings from layer 5 excitatory pyramidal neurons in the somatosensory cortex at P9, P14, and P60 in APC cKO and WT mice and quantified synaptic inhibition. Consistent with published reports (Luhmann and Prince, 1991; Le Magueresse and Monyer, 2013), the frequency of synaptic inhibition was very low at young ages and increased over time in both APC cKOs and WT mice. Interestingly, changes in synaptic inhibition were complex in APC cKOs with developmentally specific changes in IPSC frequency and amplitude. At P9, the cumulative distribution of sIPSC amplitude was significantly right shifted in APC cKOs, consistent with larger sIPSCs (Fig. 9a,d), but the frequency of sIPSCs was normal (cKO 2.03 ± 0.75 Hz; WT 1.20 ± 0.36; Fig. 9a,d). At P14, the mean sIPSC frequency was significantly decreased (cKO 5.77 ± 0.77 Hz; WT 8.86 ± 1.26 Hz; Fig. 9b,e) and the cumulative distribution of sIPSC inter-event interval was significantly right shifted in APC cKO mice (Fig. 9e), consistent with less frequent sIPSCs. At P14, the amplitude of sIPSCs was normal (cKO 18.6 ± 2.04 pA; WT 20.3 ± 1.46 pA; Fig. 9b,e). At P60, the mean sIPSC amplitude was significantly larger (cKO 21.3 ± 1.60 pA; WT 13.0 ± 0.74 pA; Fig. 9c,f) and cumulative distribution was significantly right shifted, consistent with larger sIPSCs while the frequency of sIPSC was unaltered (Fig. 9f).

**Fig 9.**
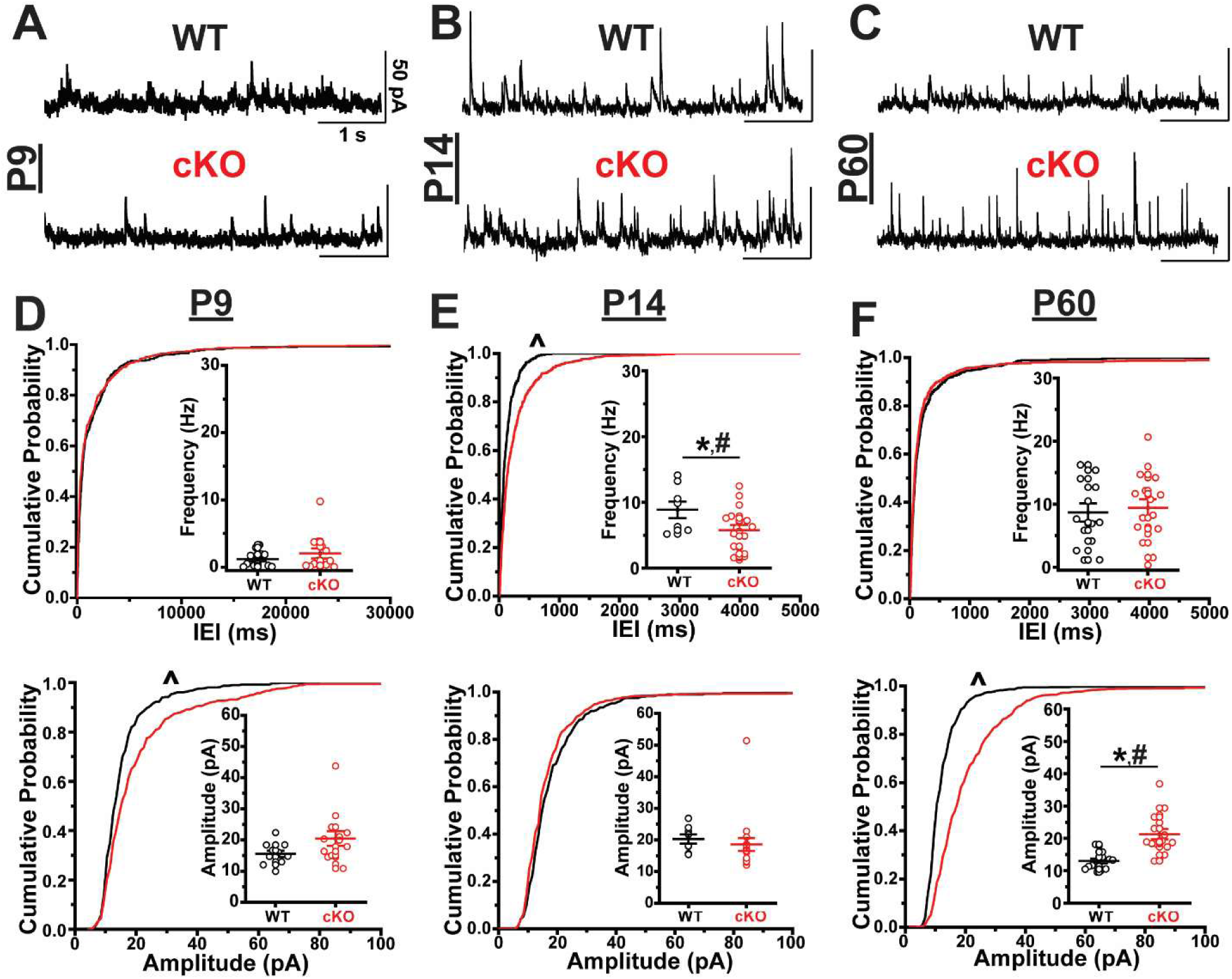
`Inhibitory spontaneous output is disrupted in the cortex of APC cKO mice.` Example spontaneous inhibitory post-synaptic current (sIPSC) traces from voltage-clamped, L5 pyramidal neurons held at 0 mV at `A.` P9, `B.` P14, and `C.` P60 for WT (black) and cKO (red) mice. All recordings made in somatosensory cortex. Scale bars = 50 pA, 1 sec. `D.` Cumulative probability plots of inter-event intervals (p=0.02, K-S test) with inset of mean frequency for WT (black, n=11 cells, 3 animals) and cKO (red, n=13 cells, 4 animals; p=0.45, U=58, Wilcoxon rank-sum test; p=0.24, t(19)=2.57, LMM) and amplitude (p=3.43e-7, K-S test) with inset of mean amplitude for WT (n=11 cells, 3 animals) and cKO (n=13 cells, 4 animals; p=0.11, U=43, Wilcoxon rank-sum test; p=0.07, t(24)=-1.88, LMM) at P9. `E.` Cumulative probability plots of inter-event intervals (p=2.29e-11, K-S test) with inset of mean frequency for WT (n=8 cells, 3 animals) and cKO (n=18 cells, 4 animals; p=0.04, t(24)=2.16, two sample t-test; p=0.03, t(21)=2.41, LMM) and amplitude (p=0.04, K-S test) with inset of mean amplitude for WT (n=8 cells, 3 animals) and cKO (n=18 cells, 4 animals; p=0.60, U=44, Wilcoxon rank-sum test; p=0.60, t(21)=0.55, LMM) at P14. `F.` Cumulative probability plots of inter-event intervals (p=0.21, K-S test) with inset of mean frequency for WT (14 cells, 4 animals) and cKO (16 cells, 6 animals; p=0.71, t(28)=0.38, two sample t-test; p=0.85, t(25)=-0.19, LMM) and amplitude (p= 4.84e-47, K-S test) with inset of mean amplitude for WT (n=14 cells, 4 animals) and cKO (n=16 cells, 6 animals; p=1.03e-4, t(28)=4.52, two sample t-test; p=5.38e-3, t(25)=-3.51, LMM) at P60. ^p<0.00001, KS test. *p<0.05, two-sample t test. #p<0.05, LMM, effect of genotype. Each data point represents one cell. Data for inset graphs are mean ± SEM.

When quantifying mIPSCs, we found that mean mIPSC frequency was not changed at any age (Fig. 10), but the cumulative distribution of mIPSC inter-event interval was left shifted in P9 APC cKOs, compared to WT, which appears to be driven by the distribution of cKO mice having less non-zero values (Fig. 10a,d). At P14 there was no difference in mIPSC frequency (cKO 4.31 ± 1.01 Hz; WT 4.00 ± 0.85 Hz; Fig. 10b,e) or amplitude (cKO 16.3 ± 0.94 pA; WT 17.3 ± 1.18 pA; Fig. 10b,e) between APC cKO and WT mice. At P60, the mean mIPSC amplitude was significantly increased (cKO 21.6 ± 2.10 pA; WT 12.7 ± 0.98 pA; Fig. 9c,f) and the cumulative distribution of mIPSC amplitude was right shifted (Fig. 10f), consistent with larger mIPSCs. Interestingly, mIPSC amplitude in APC cKO mice was significantly affected by sex. Female APC cKO mice had significantly larger mIPSCs (24.9 ± 2.16 pA) compared to APC cKO males (16.2 ± 1.46 pA; p=0.03, two sample t-test; p=9.62e-4, LMM).

**Fig 10.**
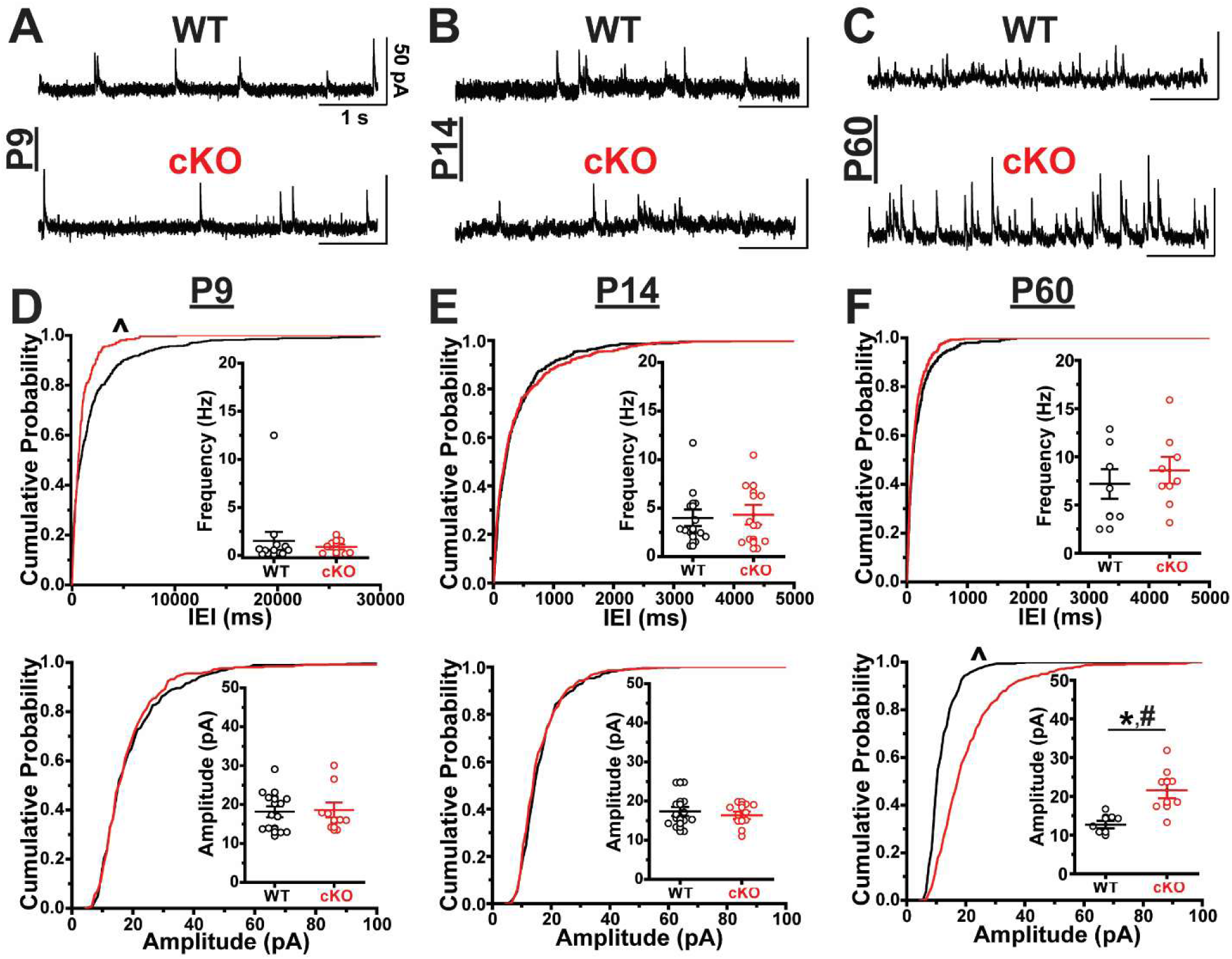
`Inhibitory miniature output is disrupted in the cortex of APC cKO mice.` Example miniature inhibitory post-synaptic current (mIPSC) traces from voltage-clamped, L5 pyramidal neurons held at 0 mV at `A.` P9, `B.` P14, and `C.` P60 for WT (black) and cKO (red) mice. All recordings made in somatosensory cortex. Scale bars = 50 pA, 1 sec. `D.` Cumulative probability plots of inter-event intervals (p=6.41e-7, K-S test) with inset of mean frequency for WT (black, n=13 cells, 3 animals) and cKO (red, n=8 cells, 3 animals; p=0.54, U=43, Wilcoxon rank-sum test; p=0.45, t(16)=0.83, LMM) and amplitude (p=0.73, K-S test) with inset of mean amplitude for WT (n=13 cells, 3 animals) and cKO (n=9 cells, 3 animals; p=0.84, U=55, Wilcoxon rank-sum test; p=0.30, t(17)=-1.09, LMM) at P9. `E.` Cumulative probability plots of inter-event intervals (p=0.11, K-S test) with inset of mean frequency for WT (n=12 cells, 3 animals) and cKO (n=10 cells, 3 animals; p=0.92, U=58, Wilcoxon rank-sum test; p=0.93, t(17)=-0.09, LMM) and amplitude (p=0.20, K-S test) with inset of mean amplitude for WT (n=12 cells, 3 animals) and cKO (n=10 cells, 3 animals; p=0.53, t(20)=0.65, two sample t-test; p=0.64, t(17)=0.50, LMM) at P14. `F.` Cumulative probability plots of inter-event intervals (p=0.10, K-S test) with inset of mean frequency for WT (n=7 cells, 3 animals) and cKO (n=8 cells, 4 animals; p=0.51, t(13)=0.68, two sample t-test; p=0.48, t(10)=-0.73, LMM) and amplitude (p=2.52e-31, K-S test) with inset of mean amplitude for WT (n=7 cells, 3 animals) and cKO (n=8 cells, 4 animals; p=2.85e-3, t(13)=3.67, two sample t-test; p=0.03, t(10)=-2.79, LMM) at P60. ^p<0.00001, KS test. *p<0.05, two-sample t test. #p<0.05, LMM, effect of genotype. Each data point represents one cell. Data for inset graphs are mean ± SEM.

In conclusion, L5Ps receive disrupted inhibitory synaptic input at all ages we examined. Most significantly, inhibitory synaptic transmission is increased onto L5Ps at P9, decreased at P14, and again increased at P60. We suspect that this is driven by premature increases in synaptic excitation of PV+INs at P9, leading to increased inhibitory output, followed by compensatory changes that result in decreased inhibition at P14. In P60 animals, increased s/mIPSC amplitude suggest further compensation, perhaps in response to abnormal synaptic excitation and seizures known to exist in APC cKOs. Increases in mISPC amplitude were sex-specific in APC cKOs.

### Intrinsic Excitability of PV+ Interneurons is Reduced in APC cKO Mice

Finally, to ascertain if altered PV+IN maturation in APC cKO mice changed the intrinsic electrophysiological and action potential firing properties of these cells, we performed current clamp recordings from G42-GFP+ interneurons in layer 5 SSC at P9, P14, and P60 (Fig. 11). Consistent with published findings, these properties changed over time, reflecting the normal development of PV+INs (Pangratz-Fuehrer and Hestrin, 2011). Resting membrane potential, membrane resistance, rheobase, and capacitance were similar in WT and cKO mice at all ages examined (Fig. 11c,d,e). Interestingly, PV+INs fired fewer action potentials in response to depolarizing current injections at all ages in APC cKO mice, as compared to WT (Fig. 11b). This suggests that while the basic electrophysiological properties of PV+INs are not altered in APC cKOs, their ability to generate robust action potential firing in response to excitatory input is reduced.

**Fig 11.**
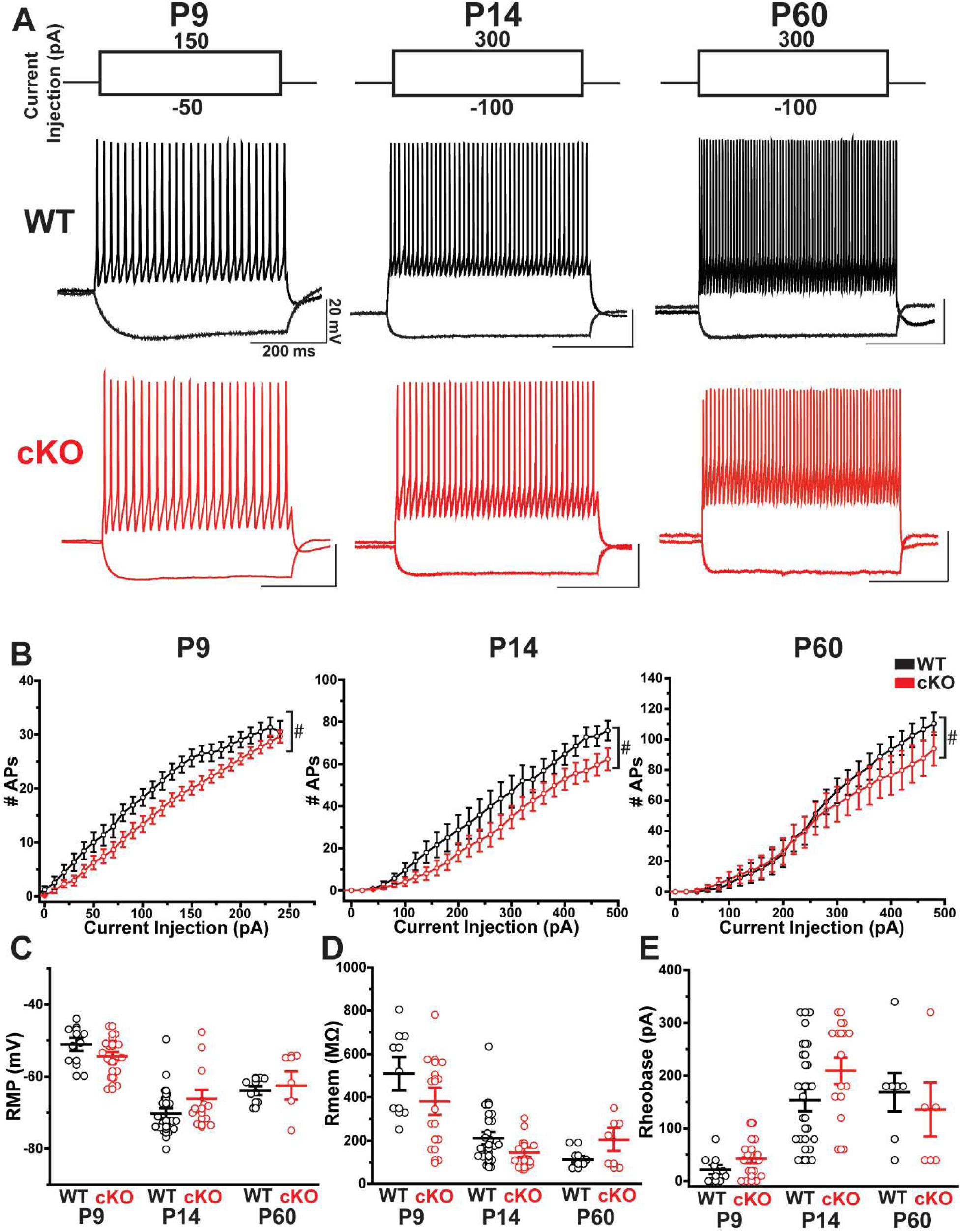
`G42-GFP+ INs are less intrinsically excitable in the APC cKO. A.` Example current clamp recordings from G42-GFP+ L5 INs during hyperpolarizing and depolarizing current injections for P9, P14, and P60 in WT (black) and cKO (red) mice. All recordings made in somatosensory cortex. Scale bars = 20 mV, 200 ms scale. `B.` Graphs of the average number of action potentials (APs) generated per current injection step for WT (black) and cKO (red) mice at P9 (p=2.25e-15, t(1014)=8.06; LMM), P14 (p<2.00e-16, t(2088)=12.2; LMM) and P60 (p<2.00e-16, t(2922)=8.37; LMM). #p<0.05, LMM, effect of genotype. Error bars indicate SEM. `C.` Resting membrane potential (mV) between WT (black) and cKO (red) mice across ages P9, P14, and P60. `D.` Membrane resistance (MΩ) between WT (black) and cKO (red) mice across ages P9, P14, and P60. `E.` Rheobase (pA) between WT (black) and cKO (red) across ages P9, P14, and P60. Each data point represents one cell. Data are mean ± SEM.

## Discussion

In this study, we examined whether the development and mature function of inhibitory GABAergic interneurons in the SSC was disrupted in the APC cKO mouse model of ISS (Summary Figure 12). Multiple ISS risk genes interact with APC and β-catenin and are important to ventral forebrain development and synaptic function (Paciorkowski et al., 2011; Pirone et al., 2017). GABAergic circuit dysfunction is thought to be a common mechanism in multiple rodent models of ISS (Marsh et al., 2009; Price et al., 2009; Olivetti et al., 2014; Marsh et al., 2016; Katsarou et al., 2018), may play a role in the human disorder (Bonneau et al., 2002; Kato and Dobyns, 2005), and has been reported in the adult PFC of APC cKO mice (Pirone et al., 2017). We found a transient reduction in immature PV+INs at P9, the peak age of behavioral spasms in APC cKO mice, using two different genetic reporter strategies (Figs. 1 and 4). We also found that in APC cKOs, the proportion of Cl-Casp3+/G42-GFP+ INs was increased at P9 but decreased at P14; the total number of Cl-Casp3+ G42-GFP+ INs was also decreased at P14 (Fig. 5). This suggests that apoptosis of developing PV+INs occurs earlier in development in APC cKO than in WT animals. Whether this affects long-term circuit function or contributes to ISS-relevant phenotypes in APC cKOs is unknown, but the role of synaptic excitation of PV+IN excitation in their maturation is well documented.

**Fig. 12.**
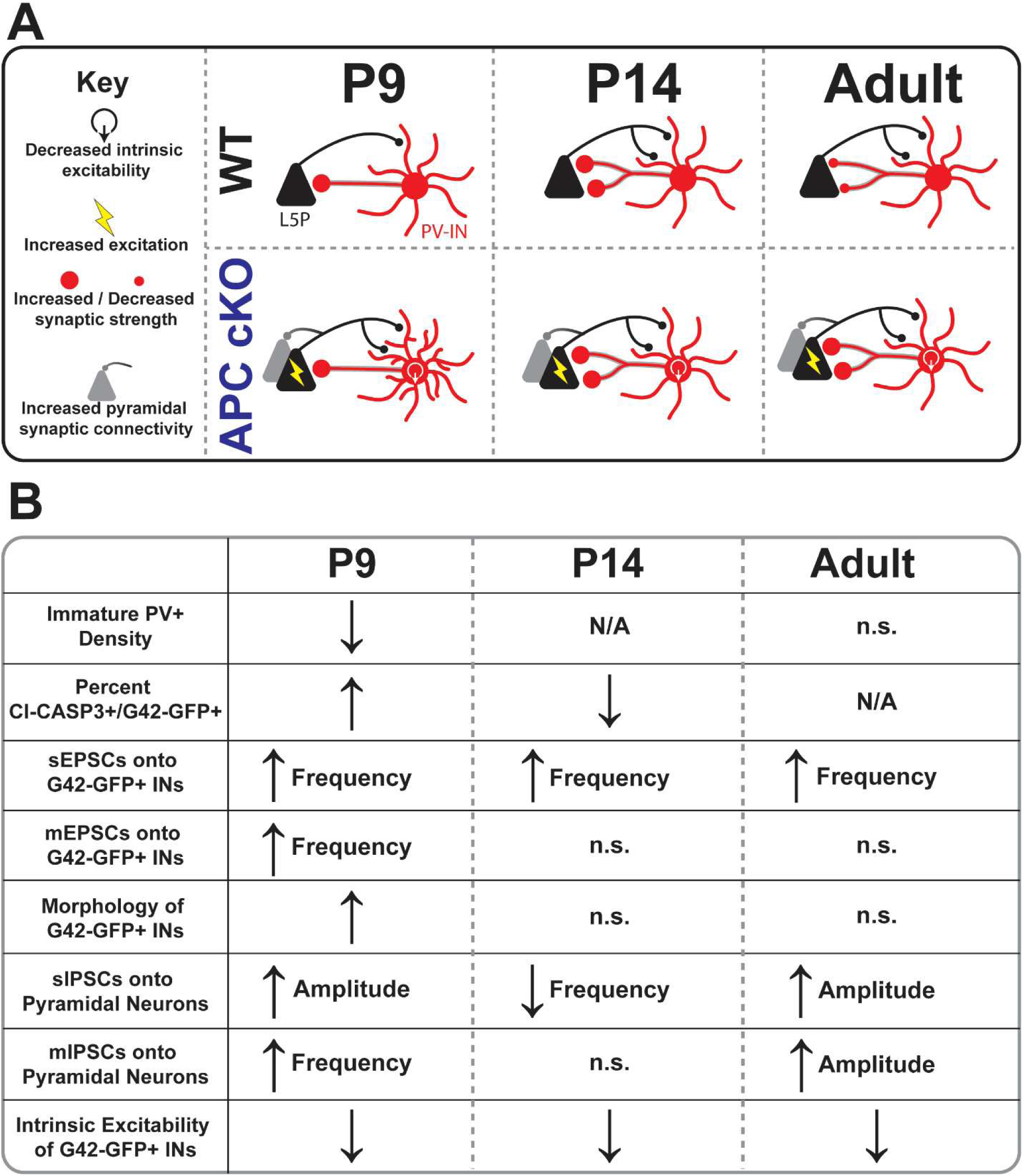
`Model of synaptic and cellular changes in APC cKO somatosensory cortical inhibitory networks. A.` Cartoon depicting synaptic changes seen in APC cKO mice. `B.` Table describing all cellular and synaptic changes seen in APC cKO mice. For A. and B., only statistically significant changes are included. For synaptic analysis only changes that were statistically significant by LMM analysis of means and KS analysis of cumulative distributions were included.

At all ages examined, PV+IN receive increased spontaneous excitatory synaptic input, but have decreased intrinsic excitability. At P9, PV+INs also have more elaborate dendritic morphology and receive more miniature excitatory synaptic input. Together, this suggests that enhanced excitatory-to-excitatory synaptic input, known to exist in APC cKO mice (Mohn et al., 2014b; Pirone et al., 2017; Pirone et al., 2018), drives aberrant circuit-mediated synaptic excitation of PV+IN. Only at P9, however, do PV+INs appear to have more excitatory synapses per neuron (suggested by increased mEPSC frequency), likely due to their increased dendritic arborization providing more area from excitatory synapses to form. Because PV+INs are less intrinsically excitable, however, this increased synaptic excitation does simply increase inhibitory GABAergic output and inhibition of nearby L5Ps in the SSC. At P9, the age of behavioral spasms, inhibition is slightly increased (increased sIPSC amplitude and mIPSC frequency, Figs. 9, 10). But at P14, inhibition is significantly decreased (lower sIPSC frequency). In adult APC cKO mice, inhibition is again heightened (increased m/sIPSC amplitude), suggesting changes in the abundance and/or composition of GABA receptors on pyramidal neurons. GABA receptor expression is highly dynamic (Kittler and Moss, 2003; Chen et al., 2012), can be regulated by neuronal activity, and is often altered in epilepsy (Brooks-Kayal and Russek, 2012). Changes in GABA receptor composition on pyramidal neurons in the SSC may also reflect compensatory changes in response to elevated excitation in APC cKOs (Gaiarsa et al., 2002). This change may also be sex-specific, as mIPSC amplitude, but not sIPSC amplitude, was increased in female APC cKOs, as compared to males. The direct relationship between these changes in GABAergic transmission and ISS-related phenotypes is yet to be determined, but it is clear that inhibitory network maturation and function are altered in APC cKO mice.

Because activity of GABAergic INs plays such an important role in their maturation (Lin et al., 2008; Huang, 2009; De Marco Garcia et al., 2011; Flores and Mendez, 2014), aberrant excitation of PV+INs in APC cKOs could have important long-term implications on inhibitory systems. Neuronal activity regulates PV+IN developmental apoptosis, with increased excitation of PV+INs favoring cell survival (Southwell et al., 2012; Priya et al., 2018; Wong et al., 2018). Surprisingly, developing PV+INs receive more excitatory synaptic input in APC cKOs (Fig. 9, 10), but appear to undergo earlier cell death (Fig. 5). We also show that PV+INs are less intrinsically excitable at all ages. Even though PV+IN receive more synaptic exaction, they may not be more active in terms of action potential firing. Because cell survival during developmental apoptosis appears to be calcium-dependent (Southwell et al., 2012; Priya et al., 2018; Wong et al., 2018), PV+IN may undergo early apoptosis in APC cKO mice because increased synaptic excitation fails to drive increased action potential generation. Of note, at P9 the total number of Cl-Casp3+/G42-GFP is unchanged, but the ratio of G42-GFP+ IN that are Cl-Casp3+ is significantly increased. This suggests there could be alterations in the generation and migration of GABAergic INs and that the ratio of Cl-Casp3+/G42-GFP+ is increased in APC cKOs due to fewer cells migrating to the cortex. A few factors argue against this possibility. First, multiple studies show that CamKIIα-Cre is not active embryonically, when GABAergic INs are generated and migrate to the cortex, making it unlikely that either process would be disrupted in APC cKOs. Second, G42-GFP+ IN density is reduced in SSC, PFC, and motor cortex at P9 (Fig. 3) eliminating the possibility that fewer G42- GFP+ INs are found in SSC because they migrate to other cortical regions. Finally, there is no change in SST+ IN density in APC cKO SSC (Fig. 2), suggesting that there are not broad changes in IN migration and number, but rather subtype-specific changes in the early development of GABAergic systems, even among MGE-derived INs.

In addition to undergoing programmed cell death, PV+INs undergo significant structural and functional changes during the first weeks of development. The development of inhibitory synapses and PV+IN perisomatic inhibition relies on neuronal activity (Gabbott and Stewart, 1987; Chattopadhyaya et al., 2004). The formation of feed-forward inhibitory circuits, mediated by cortical PV+INs, is also controlled by early neuronal activity and reciprocal synaptic connections between GABAergic INs (Marques-Smith et al., 2016; Tuncdemir et al., 2016). Therefore, the changes in synaptic transmission we report in APC cKOs could induce long-term changes in circuit function. PV+IN excitability is also highly dynamic during development, with action potential firing rate increasing over the first weeks of postnatal development (Pangratz-Fuehrer and Hestrin, 2011). The development of fast-spiking properties relies on Kv3.1 potassium channels (Goldberg et al., 2005; Goldberg et al., 2011) and the ion channel composition of the axon-initial segment (Goldberg et al., 2008). We suspect that the decreased intrinsic excitability of PV+INs that we report may arise from disruption of potassium channel expression in APC cKO, as V_m_ and Rm are unaltered in APC cKOs and changes in firing rates occur predominantly at high rates of activity (Fig. 11). GABAergic signaling also promotes the inhibitory circuit development (Huang, 2009; Wu et al., 2012), so changes in synaptic inhibition and PV+IN excitability could contribute to further changes in GABAergic function. Finally, because dendritogenesis relies on neuronal activity (Patz et al., 2004; Cohen et al., 2016), increased excitation of PV+INs may contribute to increased dendritic complexity in APC cKOs. Importantly, the role of PV+INs in ISS-relevant phenotypes must be considered in the light of other changes known to occur in excitatory neurons (Mohn et al., 2014a; Pirone et al., 2017) and other GABAergic interneurons in other brain regions (Pirone et al., 2018).

GABAergic INs are reduced in multiple models of ISS (Marsh et al., 2009; Price et al., 2009; Olivetti et al., 2014; Marsh et al., 2016; Katsarou et al., 2018), human ISS (Bonneau et al., 2002; Kato and Dobyns, 2005), and other forms of early life epilepsy (Pancoast et al., 2005), suggesting a link between GABAergic dysfunction and early life ISS-related pathologies in APC cKO mice. Of particular note, increased GABAergic signaling at P9 (Figs. 9, 10), an age at which GABA is excitatory (Ben-Ari, 2002; Ben-Ari et al., 2007), could contribute to abnormal EEG activity and behavioral spasms. In addition, increased s/mIPSC amplitude at P60 could drive hyper- synchronization of cortical circuits or compromise GABAergic inhibition by increasing intracellular chloride levels (Deeb et al., 2013; Cohen et al., 2016), potentially contributing to seizure initiation. It is challenging, however, to directly link any of the changes we report here with ISS-relevant phenotypes in APC cKO mice.

Studies in other models of ISS also underscore the complex relationship between GABAergic signaling and ISS. For example, the changes in GABAergic INs we report here, and in other studies of the APC cKO mice, suggest unique changes in PV+ and SST+ INs. Interestingly in other well established models of ISS, namely the multiple hit rat model (Scantlebury et al., 2010; Raffo et al., 2011; Katsarou et al., 2018), PV+INs, but not SST+ or calretinin-positive GABAergic cells, are selectively lost (Katsarou et al., 2018). In the Arx expansion model of ISS, there is a decrease in calbindin-positive and neuropeptide Y GABAergic INs, but not SST+, PV+, or calretinin+ INs (Price et al., 2009; Olivetti et al., 2014; Lee et al., 2017). Targeted deletion of Arx also results in a decreased density of calbindin, but not PV+, interneurons (Marsh et al., 2009). Interestingly, different mutations in Arx are associated with distinct changes in the density of specific GABAergic IN subpopulations (Olivetti and Noebels, 2012), adding further complexity. Of relevance to our study, there is increased apoptosis during cortical development (P7) in the Arx expansion model of ISS, although this does not occur in Arx+ cells but in a non-cell-autonomous fashion (Siehr et al., 2020). Because Arx signaling may share interactions with APC/β-catenin signaling (Cho et al., 2017; Pirone et al., 2017), it is particularly interesting that different populations of GABAergic INs are affected in each model. The molecular pathology may be completely distinct in these models, developmental differences in when dysfunction occurs (post- natal in the APC cKO and multiple-hit rat model; in utero in the Arx expansion model) may contribute to model-specific changes in GABAergic INs, or unique changes in gene expression, epigenetic state, splicing, or other developmental diversification of GABAergic INs may be affected in way we do not yet understand (Mayer et al., 2018; Wamsley et al., 2018; Allaway et al., 2021). Conversely, loss of PV+IN in both the APC cKO and multiple-hit rat model suggest changes in PV+IN may have more to do with the presence of infantile spasms, rather than a specific disruption in APC/β-catenin signaling. Clearly, future work will be required to elucidate the molecular mechanisms driving disruptions in GABAergic INs in ISS and how those changes contribute to disease pathology.

While our findings demonstrate disruptions in inhibitory network development and mature function in the APC cKO mouse, there are open questions and caveats that remain. First, identifying immature PV+INs at P9 is challenging because PV protein is not expressed until later in life. To increase rigor, we have two genetic reporter lines (G42 and Lhx6-GFP) to quantify immature PV+IN density. Novel viral approaches (Dimidschstein et al., 2016; Vormstein-Schneider et al., 2020) and advances in our molecular understanding of PV+IN maturation (Fishell, 2007; Karayannis et al., 2014; Mayer et al., 2018; Allaway et al., 2021) may enhance our ability to identify and manipulate immature PV+INs with increased confidence. Second, we do not find that decreases in PV+IN density persist into adulthood in the SSC. We suspect this may be brain region specific, as we have previously shown decreases in both PV+ and SST+ interneuron densities in the prefrontal cortex of adult cKO mice (Pirone et al., 2018). Third, there is increased variability in G42-GFP+ IN and PV+IN cell density in APC cKO mice compared to WT littermates. This is not entirely surprising as there is significant heterogeneity of behavioral spasms and seizures in APC cKOs (Pirone et al., 2017). Whether the variability in GABAergic IN density is related to heterogeneity in ISS-relevant phenotypes remains to be seen. Variability in reporter expression may also contribute as GFP+ expression in G42 cells increases over the second postnatal week (Chattopadhyaya et al., 2004), the same time window in which we measure IN densities. Again, use of the Lhx6-GFP+ reporter, which has more stable GFP expression in early development, helps mitigate this concern. Fourth, we do not address whether altered Wnt/β- catenin signaling in excitatory neurons leads to non-cell autonomous effects in nearby PV+INs, including changes in IN proliferation and/or migration. Although Wnt/β-catenin signaling affects IN proliferation and cell fate decisions (Backman et al., 2005; Gulacsi and Anderson, 2008; Paina et al., 2011; Allaway et al., 2021), those events occur before CamKIIα-Cre expression in APC cKO mice (Pirone et al., 2017). Finally, this study was not designed or powered to address sex as a biological variable and animal sex was not recorded for all experiments, especially at younger ages. Due to this, we cannot evaluate the role of sex as a biological variable for many of our findings. Interestingly, increases in mIPSC amplitude, but not sIPSC amplitude, was increased in APC cKO females compared to APC cKO males. This is consistent with sex-specific changes reported in other ISS models (Marsh et al., 2009; Ono et al., 2011; Briggs et al., 2014; Olivetti et al., 2014; Dube et al., 2015; Frost et al., 2015; Gataullina et al., 2016; Siehr et al., 2020; Akman et al., 2021; Loring et al., 2021) and indicates sex-specific changes should be investigated in future studies. No effects of sex on adult density of PV+INs, G42-GFP-INs, Lhx6-GFP+ INs, or SST+ INs, or any measures of inhibitory synaptic transmission were seen. Future studies will be powered to examine sex as a biological variable on all outcomes.

Those caveats aside, determining whether interneuron dysfunction during early development is sufficient to drive behavioral infantile spasms and later electroclinical seizures is of great scientific and clinical interest. The evidence from human and mouse models of ISS, including the APC cKO mouse, suggest a role for IN dysfunction in ISS and implicate them in the circuit mechanisms that drive both infantile spasms and adult seizures. Our work points to P9 as a pivotal developmental time point in the APC cKO mouse, with peak behavioral spasms (Pirone et al., 2017) coinciding with decreased PV+IN density, an increased proportion of cell death in G42-GFP+ INs, increased glutamatergic input, and increased dendritic complexity of PV+INs. In addition, this highlights a critical window during the second postnatal week for inhibitory circuit integration, in which feedback and feedforward inhibition is the ultimate outcome in normal development (Gabernet et al., 2005; Cruikshank et al., 2007; Lim et al., 2018). Disruption in neuronal activity, ambient glutamate levels, and interneuron maturation during this time window can lead to long-term changes in circuit structure and function (Andresen et al., 2014; Tuncdemir et al., 2016; Lau et al., 2017; Hanson et al., 2019) and may contribute to ISS-relevant phenotypes. Finally, our findings show that excitatory neuron dysfunction can initiate long-term interneuron dysfunction and further implicate GABAergic dysfunction in ISS, even when pathology is initiated in other neuronal types. They also underscore the importance of clinical efforts to identify patients with ISS very early in life for therapeutic intervention (Shields, 2006). In summary, our findings show that PV+INs and cortical inhibitory networks are disrupted during early development in APC cKO mice and suggest that this may contribute to the manifestation of spasms and later seizures.

## ACKNOWLEDGMENTS

We would like to thank members of the Dulla Lab, Jacob lab, Oliva Drake, and Elisabeth Lawton for their helpful discussions. Research was supported by R01 NS100706 (CD), R56 NS094889 (CD), CURE Epilepsy (CD and MJ), and American Epilepsy Society Predoctoral Fellowship (RR).

